# CPSF6 Promotes HIV-1 Preintegration Complex Function

**DOI:** 10.1101/2025.01.28.635394

**Authors:** Evan Chaudhuri, Sooin Jang, Rajasree Chakraborty, Rajalingam Radhakrishnan, Bjarki Arnarson, Prem Prakash, Daphne Cornish, Nicholas Rohlfes, Parmit K. Singh, Jiong Shi, Christopher Aiken, Edward Campbell, Judd Hultquist, Alan N. Engelman, Muthukumar Balsubramaniam, Chandravanu Dash

## Abstract

Cleavage and polyadenylation specificity factor 6 (CPSF6) is part of the cellular cleavage factor I mammalian (CFIm) complex that regulates mRNA processing and polyadenylation. CPSF6 also functions as a HIV-1 capsid (CA) binding host factor and promotes viral DNA integration targeting into gene dense regions of the host genome. However, the effects of CPSF6 on the activity of the HIV-1 preintegration complex (PIC) - the machinery that carries out viral DNA integration to establish infection - is unknown. To study CPSF6’s role in HIV-1 PIC function, we extracted PICs from cells depleted of CPSF6 or expressing a CPSF6 mutant that cannot bind to CA. These PICs exhibited significantly lower integration activity when compared to the control PICs. Addition of recombinant CPSF6 restored the integration activity of PICs extracted from the mutant cells, suggesting a direct role of CPSF6 in PIC function. To solidify CPSF6’s effect on PIC function, we inoculated CPSF6-depleted and CPSF6-mutant cells with HIV-1 particles and measured viral DNA integration into the host genome. A significant reduction in viral integration in these cells was detected and this defect was not a consequence of reduced reverse transcription or nuclear entry. Additionally, mutant viruses deficient in CA-CPSF6 binding showed no integration defect in CPSF6 mutant cells. Finally, sequencing analysis revealed that HIV-1 integration in the CPSF6 mutant cells was significantly redirected from the gene dense regions of the host genome. Collectively, these results suggest that CPSF6-CA interaction regulates PIC function both in vitro and in infected cells.

**IMPORTANCE:** HIV-1 infection is dependent on the interaction of the virus with host factors. However, the molecular details of virus-host factor interactions are not fully understood. For instance, HIV-1 capsid provides binding interfaces for several host factors. CPSF6 is one such capsid-binding host factor, whose cellular function is to regulate mRNA processing and polyadenylation. Initial work identified a truncated cytosolic form of CPSF6 that restricted HIV infection by blocking viral nuclear entry. However, it is now established that the full-length CPSF6 primarily promotes integration targeting into gene dense regions of the host genome. Here we report that CPSF6-CA interaction promotes the activity of HIV-1 preintegration complexes (PICs). We also observed that disruption of CPSF6-CA binding in target cells significantly reduced viral integration and directed integration targeting away from gene-dense regions. These findings demonstrate a critical role for the CPSF6-CA interaction in PIC function and integration targeting.

## INTRODUCTION

Approximately eighty-two million people have been infected with the human immunodeficiency virus 1 (HIV-1) [1]. HIV-1 infects immune cells such as T-lymphocytes, monocytes, and macrophages that express CD4 receptor and CCR5 or CXCR4 co-receptors [2–5] [6–8]. HIV-1 infection results in the steady decline of CD4+ T-cells and the development of immunodeficiency [9, 10]. HIV-1 infection-associated immunodeficiency ultimately progresses to acquired immunodeficiency syndrome (AIDS), a deadly disease that has already killed more than 42 million people worldwide. Fortunately, the development and access to highly effective antiretroviral therapy (ART) has transformed HIV-1 infection to a chronic disease and has dramatically reduced AIDS-related deaths. However, these therapies require lifelong administration, have side-effects, and face drug resistance [4]. Therefore, identification of novel therapeutic targets is necessary to effectively control HIV-1. Understanding the mechanisms by which host factors promote HIV-1 infection has the potential to identify such antiviral targets.

To begin infection, HIV-1 envelope binds to its target cell receptor (CD4+) and coreceptor, causing fusion of the viral membrane with the cellular plasma membrane and the release of the viral capsid into the cytoplasm [11]. The HIV-1 capsid contains two copies of a linear single-stranded (ss) viral RNA genome as well as viral and host factors [12–17]. The capsid shell is composed of two distinct capsomeres of the viral capsid (CA/p24) protein [18, 19], a major capsomere as a hexamer of CA; and a minor capsomere formed of a pentamer of CA [13, 20–22]. The capsid shell is made up of ∼200 CA hexamers and exactly 12 pentamers [14, 18, 23]. The fullerene curvature of the HIV-1 capsid is formed by seven pentamers at the broad end and five pentamers at the narrow end, with the rest of the capsid composed of hexamers. Although the exact mechanism of CA oligomerization is not fully understood, CA monomers are connected by specific CA-CA interactions that determine assembly and stability of the capsid. Contacts between the N-terminal domain (NTD)-NTD of CA monomers stabilize the hexamers and pentamers and form a central pore. The C-terminal domain (CTD) of CA participates in dimeric and trimeric interactions between individual hexamers and pentamers. Overall, these interactions generate at least four known interfaces for host factor binding-1) six hydrophobic clefts within each hexamer known as phenylalanine-glycine (FG) pockets, 2) the arginine (R18) pore at the center of the hexamer, 3) a cyclophilin A (CypA) binding loop on each CA-NTD subunit, and 4) an electronegative patch formed at the vertex of three adjoining hexamers [18, 19, 24–35]. Binding of host factors to these pockets of the capsid critically regulates HIV-1 infection [35].

From the host cytoplasm, the HIV-1 capsid is trafficked towards the nuclear pore complex (NPC) [36, 37]. HIV-1 nuclear import is dependent on the capsid [38–40] and is regulated by the nucleoporins (Nups) of the NPC [41]. The human NPC is constructed from ∼33 Nups arranged in an 8-fold rotational symmetry [42] and the HIV-1 capsid contains binding sites for several Nups. The large outer nucleoporin, Nup358 (a.k.a. RAN binding protein 2 or RanBP2), binds to the capsid via its cyclophilin-homology domain [19, 43, 44], which likely docks the capsid at the NPC [45–48]. Early models indicated that the HIV-1 capsid would disassemble at the NPC to allow for translocation of the viral preintegration complexes (PICs) through the central channel [49]. However, more recent work has suggested NPC plasticity can accommodate the capsid at its widest point (∼60 nm) [50]. Microscopy-based studies have also suggested that HIV-1 capsids remain largely intact during nuclear import [51, 52]. Additionally, NUP153, located at the nucleoplasmic side of the NPC, as well as the nuclear host factor cleavage and polyadenylation specificity factor 6 (CPSF6), display high affinity binding to the capsid lattice compared to monomeric CA [48, 53], suggesting that at least some portion of the capsid lattice is sustained during and after nuclear entry [51, 52, 54–57].

Recent work has helped to clarify the contributions of Nups to HIV-1 nuclear transport. About one-third of human Nups are enriched for FG dipeptide repeats and are accordingly referred to as FG-Nups. Several FG-Nups, including Nup62, Nup98, Nup58, Nup42, POM121, and Nup153, bind the HIV-1 capsid FG pocket [58–61]. CPSF6 also harbors an FG dipeptide that binds this same region of capsid [62, 63]. While en route to the nucleus, the capsid core-associated reverse transcription complex (RTC), containing the viral reverse transcriptase (RT) and integrase (IN) enzymes, synthesizes a double-stranded DNA copy of the viral RNA genome [37, 64]. IN-mediated processing of the viral DNA ends operationally morphs the RTC into PIC. The PIC then integrates the viral DNA into the host genome [65]. PICs were first described as high molecular weight complexes in cell extracts that integrated the viral DNA made by reverse transcription into an exogenous target DNA in vitro [66, 67]. Aided by host factors, the PIC integrates viral DNA preferentially into actively transcribed genes of the host genome [65]. Integration establishes a permanent provirus in the host genome that serves as the genetic element to produce progeny virions [65].

CPSF6 was first implicated in HIV-1 biology via a genetic screen for mouse-specific restriction factors [68], since mouse cells do not support HIV-1 infection [69]. This study identified a C-terminal truncated CPSF6, called CPSF6-358, which binds the HIV-1 capsid [68]. CPSF6 is conserved among mammals and is involved in mRNA processing and polyadenylation as part of the cellular pre-mRNA processing complex cleavage factor Im (CFIm) [70]. In humans, *CPSF6* is located on chromosome 12 (12q15) and encodes a predominant 551-amino acid (aa) splice variant, which is associated with nuclear paraspeckles. Structurally, CPSF6 has three key regions. Near the N-terminus (aa 81-157) is the RNA recognition motif (RRM) that binds to CPSF5 in the CFIm complex [71, 72]. The C-terminal (aa 489-551) arginine/serine-like domain (RSLD) engages the β-karyopherin transportin 3 (TNPO3; a.k.a. TRN-SR2) and acts as the nuclear localization signal for CPSF6 [73, 74]. The RSLD also displays liquid-liquid phase separation (LLPS) activity in vitro [75] and confers higher-order capsid binding, which is critical for HIV-1 capsid to move downstream from the nuclear rim during infection [76]. The central region of CPSF6 (aa 208-398) harbors a proline-rich domain (PRD) with a prion-like low complexity region (aa 217-326), including the FG dipeptide that directly engages with the HIV-1 capsid [53, 63].

The role of CPSF6 in HIV-1 replication is multipronged. Initially, CPSF6-358 lacking the RSLD was reported to inhibit HIV-1 nuclear import [68]. Passaging experiments in the presence of CPSF6-358 produced viruses bearing an N74D substitution in CA, which was resistant to CPSF6-358 mediated inhibition of HIV-1 [68, 77]. These results indicated that CA was the viral determinant for CPSF6-358 restriction [68, 77]. A separate mutation near the N74 residue, A77V, was also reported to reduce CPSF6 interaction with the HIV-1 CA [78]. CPSF6 depletion can marginally inhibit HIV-1 nuclear import [79] and, in monocyte/macrophages, CPSF6 protects HIV-1 replication complexes from innate immune sensing [80, 81]. However, CPSF6 appears to primarily regulate intranuclear localization of HIV-1 PICs for integration site selection [79, 82, 83]. For instance, CPSF6 enables trafficking of HIV-1 cores to colocalize with nuclear speckles for subsequent PIC-mediated integration into speckle-associated domains (SPADs) of human DNA [76, 84]. CPSF6 also seems to coordinate with other capsid-binding host factors, such as CypA, to orchestrate the early steps of HIV-1 infection [85]. The mechanism by which CPSF6 regulates PIC function has not been previously studied.

Here, we used human cell lines depleted of CPSF6 or expressing a CPSF6 mutant (FG>AA) that cannot bind to HIV-1 CA [53] for PIC-based experiments. Such PICs exhibited significantly lower integration activity compared to the PICs from the control cells, whereas addition of purified CPSF6 protein stimulated PIC activity. To study the effects of CPSF6 on PIC function during HIV-1 infection, we quantified DNA replication intermediates (reverse transcription, 2-long terminal repeat (LTR) circles for nuclear entry, and integration) and evaluated integration site selection profiles. Our results showed that disrupting CPSF6-CA binding reduced HIV-1 integration without measurably reducing reverse transcription or nuclear entry. Further, the CPSF6-CA binding deficient mutant viruses N74D and A77V showed minimal integration defects in the CPSF6 mutant cells. We also found that disruption of CPSF6-CA binding significantly retargeted viral DNA integration into lamina-associated domains (LADs) instead of the gene-dense SPADs. These results for the first time support a direct role for CPSF6 in PIC function.

## MATERIALS AND METHODS

### Chemicals, plasmids and other reagents

The antiretroviral compound Raltegravir was obtained through BEI Resources (NIH HIV Reagents Program; Manassas, VA) and prepared as previously described [86]. The viruses utilized in this study were generated from the HIV-1 molecular clone pNLX.Luc(R-)ΔAvrII [87], and its CA mutant derivatives CA-N74D and CA-A77V [87]. The plasmid pHCMV-G encoding VSV-G [88] was used in co-transfection experiments for generating pseudotyped HIV-1 particles. Recombinant CPSF6 protein expressed in *E. coli* strain BL21(DE3) was purified as previously described [76].

### Western blot

To detect protein levels by western blot, cell lysates were prepared using 1X RIPA buffer (Sigma-Aldrich) and total protein concentrations were quantified by BCA protein assay (Pierce). Equal amounts of total protein from cell lysates were electrophoresed on 4-12% sodium dodecyl sulfate (SDS)-polyacrylamide gels and transferred to nitrocellulose membranes using a semidry blotter (Bio-Rad). Membranes were blocked with 5% (w/v) nonfat milk in Tris-buffered saline with Tween 20 (TBST; pH 8.0; Chem Cruz). Blots were then probed with the primary antibody in blocking buffer (anti-CPSF6: Proteintech catalog # 15489-1-AP at 1:4,000 dilution or Abcam ab175237 at 1:10,000; anti-actin: Sigma-Aldrich catalog # A2228 at 1:60,000 dilution or A3854-200UL at 1:20,000) and subsequently probed with a secondary antibody conjugated to horseradish peroxidase (anti-rabbit: BioRad catalog # 1706515 at 1:6,000 dilution or Dako P0448 at 1:10,000; BioRad catalog # 1706516 anti-mouse at 1:40,000 dilution). All blots were washed in TBST and developed by using the enhanced chemiluminescence (ECL) procedure (Bio-Rad). Densitometry analysis was performed by using LI-COR Image Studio Digits version 5.2 software (LI-COR). Data were normalized to levels of β-actin.

### Cell culture

TZM-bl, human embryonic kidney 293T (HEK293T), HeLa, Jurkat T-cells and SupT1 cell lines were obtained from the American Type Cell Culture Collection (Manassas, VA). The TZM-bl and HEK293T cells were cultured in Dulbecco’s Modified Eagle’s medium (ThermoFisher Scientific) containing 10% heat-inactivated fetal bovine serum (FBS), penicillin (50 IU/mL), streptomycin (50 µg/mL). Jurkat and SupT1 cells were cultured in Roswell Park Memorial Institute (RPMI) 1640 medium (ThermoFisher Scientific) containing 10% heat-inactivated FBS, penicillin (50 IU/mL), streptomycin (50 µg/mL). All cell lines were cultured at 37°C with 5% CO_2_.

### CRISPR-Cas9 knock-in and knockout

Jurkat cells were genetically modified using a CRISPR-Cas9 based knockout strategy [89, 90]. For synthesis of CRISPR-Cas9 ribonucleoprotein complexes (crRNPs), lyophilized crRNA and tracrRNA (Dharmacon) were resuspended at 160 µM in a buffer containing 10 mM Tris-HCl (pH 7.4) and 150 mM KCl. 5 µL of 160 µM crRNA was mixed with 5 µL of 160 uM tracrRNA, followed by a 30-min incubation at 37°C. The crRNA:tracrRNA complexes were then gently mixed with 10 µL of 40 µM *S. pyogenes* Cas9 (UC-Berkeley Macrolab) and incubated at 37°C for 15 min. crRNPs were aliquoted into five sets of 3.5 µL each and stored at-80°C prior to use. crRNAs were obtained from the Dharmacon predesigned Edit-R library or synthesized as custom sequences (CPSF6 guide #5 GGACCACATAGACATTTACG, CPSF6 guide #6: ATATATTGGAAATCTAACAT, Non-Targeting guide #4 (U-007504-20)). To generate polyclonal CPSF6 knock-out Jurkat cells, 3 x 10^5^ cells per electroporation reaction were spun at 400xg for 5 min and supernatant was removed by aspiration. Cells were resuspended in buffer consisting of 16.4 µL SE Nucleofector solution with 3.6 µL supplement per reaction. 20 µL of cell suspension was mixed with 3.5 µL of each crRNP and transferred to 96-well electroporation cuvettes for electroporation with the 4D Core Unit using pulse code CM-137 (Lonza). 100 µL fresh RPMI media was added to each well following electroporation, and cuvettes were placed in cell culture incubator to recover for 30 min. Cells were moved to 48-well tissue culture plates prefilled with 400 µL RPMI, then incubated in a tissue culture incubator. Subsequently, single-cell clones from the gene-knockout Jurkat cell pools was isolated by limiting serial dilution. Cells were initially cultured to a density of 10^6^ cells/mL. Cells (2×10^3^) were added to the top-upper leftmost corner of a 96-well plate in 200 µL of RPMI. Cells were then serially diluted within the plate, initially by column and then by row. Plates were allowed to grow in a tissue culture incubator for 3 weeks. Subsequently, 10 clones were selected from NT control and CPSF6 knock-out Jurkat cells and transferred to 48-well plates with 200 µL RPMI added. Cell expansion was continued in 48-well plates and then further extended to T12.5 flasks as the cells were passaged. Cells were passaged around confluency of 80%. Cells from proposed monoclonal populations were then sequenced for exon 7 of *CPSF6* and were further probed for CPSF6 protein expression.

HEK293T and HeLa CPSF6 knockout (CKO) cells were previously described [53, 83, 91, 92]. To mutate the portion of CPSF6 that encodes for the capsid binding FG motif (FG>AA), a single guide RNA (sgRNA) 5’-CCCAATGGAGGCTGCCCAAA-3’ targeting exon 7 of *CPSF6* and a single-stranded oligodeoxynucleotide (ssODN) donor template 5’-GCCCTCCACCACCAGTTCTTTTTCCTGGACAACCTGCTGCGCAGCCTCCATTGGGTCCACTTCCTCCTGGCCCTC-3’ were purchased from Horizon Discovery. To form crRNPs, 5 μL of 60 μM CPSF6 sgRNA and 5 μL of 30 μM TrueCut Cas9 v2 (Thermo Fisher) were mixed and incubated at 37°C for 15 min. Approximately 1×10^6^ SupT1 cells were transfected with 10 μL of 15 μM crRNPs and 10 μL of 100 μM ssODN using the nucleofection kit and Nucleofector I device (Lonza), following the manufacturer’s protocol. Three to five days post-transfection, single cells were isolated by limiting dilution cloning in 96-well plates, and then transferred to 24-well plates as the cells propagated. To confirm biallelic knock-in, genomic DNA was extracted, and the targeted locus was amplified using primers 5’-TGGCACGAATTCGTAAGGATATACTTCATTGTAGTTGGTAGTG-3’ and 5’-CACTGAGGATCCGCGTTCTTGCAGTATCCATTTCC-3’. Sanger sequencing of PCR amplicons using primer 5’-CCATAGTCACCCCTATCATATGG-3’ was performed before and after cell cloning.

### Viral stock preparation

The viruses utilized in this study were generated from HIV-1 molecular clone pNLX.Luc(R-)ΔAvrII (wildtype (WT) and CA mutant derivatives (N74D and A77V). Infectivity and viral DNA quantification experiments were performed with VSV-G pseudotyped NL4-3 virions that were generated from the pNLX.Luc(R-)ΔAvrII plasmid that contains a frameshift in the *env* gene and the firefly luciferase-coding sequence inserted in place of the *nef* gene [87]. All virus stocks were prepared as previously described [86]. Briefly, 5 x 10^6^ HEK293T cells were seeded in 10 cm cell culture plates and cultured overnight. Next day, cells in each culture plate were co-transfected with 10 µg of pNLX.Luc(R-)ΔAvrII and pHCMV-G plasmid DNA complexed with 50 μL PEI reagent (1 mg/mL) (Sigma-Aldrich catalog # 764604), and the culture media was changed 16 h post-transfection. Forty-eight hours later, the virus-containing culture media was collected, subjected to low-speed centrifugation, and the supernatant was passed through 0.45 µm filter. The filtrate was treated with DNase I (ThermoFisher Scientific catalog #AM2238) at a final concentration of 20 μg/mL at 37°C. The viral stocks were stored in-80°C freezer until assayed for infectivity. Subsequently, ELISA was used to assay the virus preparations for p24 amounts using anti-p24 antibody as previously described [86]. Virus infectivity was determined using TZM-bl indicator cells, as described previously [93].

### Single-cycle infection assay

1×10^6^ SupT1 cells in each well of a 12-well plate flat-bottom cell culture plate were spinoculated (480 x *g*) with 450 ng of pseudotyped HIV-1.Luc for 2 h at 25°C and were then cultured for 24 h in 37°C/5% CO_2_ incubator. Subsequently, cells were pelleted (500 x *g*) and washed three times with phosphate buffered saline (PBS, pH 7.4, Thermo Fisher Scientific). Cell samples resuspended in PBS were aliquoted for assessing infectivity via luciferase assay and for total DNA isolation. For luciferase assay, cell pellet was lysed in 100 μL 1X Glo Lysis Buffer (Promega Catalog# E2661), rocked for 1 h at 25°C, and centrifuged (16000 x *g)* for 5 min to pellet cellular debris. Triplicate 25 μL aliquots of each sample supernatant were dispensed into 96-well black flat bottom plates. Luciferase activity was determined using Luciferase assay system (Promega) in a microplate reader (Biotek). Infectivity assays for HEK293T cells were performed by seeding 1×10^6^ cells in each well of a 12-well flat-bottom cell culture plate 24 h prior to addition of pseudotyped HIV-1.Luc. Cells were incubated with viral particles and spinoculated for 2 h at 25°C prior to incubation in a 37°C/5% CO_2_ incubator for 24 h. Cells were then harvested by detachment from the plate and pelleted by centrifugation (500 x g). Cell samples were processed for luciferase assay as described above for SupT1 cells.

### Total DNA isolation for qPCR assays

To isolate total DNA from infected cells, 1×10^6^ SupT1 cells in each well of a 12-well plate flat-bottom cell culture plate were spinoculated (480 x *g*) with 450 ng of pseudotyped HIV-1.Luc for 2 h at 25°C and were then cultured for 24 h in 37°C/5% CO_2_ incubator. Half of the cells were used for assessing virus infectivity via a luciferase reporter assay, and total DNA was isolated from the remaining cells by using the Quick-DNA miniprep kit (Zymo Research).

### qPCR for measuring reverse transcription and 2-LTR circles

Late-stage reverse transcription products were quantified by using SYBR green-based qPCR, and 2-LTR circles were quantified by using TaqMan probe-based qPCR. The SYBR green-based qPCR mix contained 1X iTaq Universal SYBR Green Supermix (Bio-Rad), 300 nM concentrations (each) of forward primer (5’-TGTGTGCCCGTCTGTTGTGT-3’) and reverse primer (5′-GAGTCCTGCGTCGAGAGAGC-3’), and 100 ng of total DNA from infected cells. The TaqMan probe-based qPCR mix contained 1X iTaq Universal Probe Supermix (Bio-Rad), 300 nM (each) of forward primer (5’-AACTAGGGAACCCACTGCTTAAG-3’) and reverse primer (5’-TCCACAGATCAAGGATATCTTGTC-3’), 100 nM of the TaqMan probe (5’-[FAM] ACACTACTTGAAGCACTCAAGGCAAGCTTT-[TAMRA]-3’), and 100 ng of total DNA from infected cells. The cycling conditions for SYBR green-based qPCR included an initial incubation at 95°C for 3 min, followed by 39 cycles of amplification and acquisition at 94°C for 15 s, 58°C for 30 s, and 72°C for 30 s. For the SYBR green-based qPCR, the thermal profile for melting-curve analysis was obtained by holding the sample at 65°C for 31 s, followed by a linear ramp in temperature from 65 to 95°C with a ramp rate of 0.5°C/s and acquisition at 0.5°C intervals. The TaqMan probe-based qPCR included an initial incubation at 95°C for 3 min, followed by 39 cycles of amplification and acquisition at 94°C for 15 s, 58°C for 30 s, and 72°C for 30 s. During qPCR of the samples, a standard curve was generated in parallel and under same conditions using 10-fold serial dilutions of known copy numbers (10^0^ to 10^8^) of the pNLX.Luc(R-)ΔAvrII plasmid (for late reverse transcription products) or the p2LTR plasmid containing the 2-LTR junction sequence (for 2-LTR circles). CFX Manager software (Bio-Rad) was used to analyze the data and determine the copy numbers of the late reverse transcription products and the 2-LTR circles by plotting the data against the respective standard curves.

### Alu-gag nested qPCR for measuring HIV-1 proviral integration

To measure integrated HIV-1 proviral DNA, a nested PCR method involving a first-round endpoint PCR with primers designed to amplify the junctions of chromosomally integrated viral DNA with host DNA, followed by a second round of qPCR with primers that amplify viral LTR-specific sequences present in the first round PCR amplicons, was used with some modifications. Briefly, the first-round PCR was performed in a final volume of 50 μL containing 100 ng of total DNA from infected cells, 1X Phusion reaction buffer (NEB), deoxy-nucleotide triphosphate (dNTP) nucleotide mix containing 200 µM concentrations of each nucleotide (Promega), 500 nM (each) of primers that target the host chromosomal Alu repeat sequence (5’-GCCTCCCAAAGTGCTGGGATTACAG-3’) and HIV-1 Gag sequence (5’-GTTCCTGCTATGTCACTTCC-3’), and 1.25 U of Phusion DNA Polymerase (NEB). The thermocycling conditions included an initial incubation at 95°C for 5 min; followed by 23 cycles of amplification at 94°C for 30 s, 50°C for 30 s, 72°C for 4 min and a final incubation at 72°C for 10 min. The second-round qPCR mix contained a 1/10 volume of the first-round PCR products as the template DNA, 1X iTaq Universal Probe Supermix (Bio-Rad), 300 nM (each) of the viral LTR-specific primers that target the R region (5’-TCTGGCTAACTAGGGAACCCA-3’) and the U5 region (5’-CTGACTAAAAGGGTCTGAGG-3’), and a 100 nM concentration of TaqMan probe (5’-[6-FAM]-TTAAGCCTCAATAAAGCTTGCCTTGAGTGC-[TAMRA]-3’). The qPCR cycling conditions included an initial incubation at 95°C for 3 min, followed by 39 cycles of amplification and acquisition at 94°C for 15 s, 58°C for 30 s, and 72°C for 30 s. During the second round qPCR of the samples, a standard curve was generated in parallel and under same conditions using 10-fold serial dilutions of known copy numbers (10^0^ to 10^8^) of the of the pNLX.Luc(R-)ΔAvrII plasmid. Data were analyzed using CFX Manager software (Bio-Rad), and integrated viral DNA copy numbers were determined by plotting the qPCR data against the standard curve.

### Extraction of HIV-1 PICs

HIV-1 PICs were extracted from infected T-cells and HEK293T using a modified version of previously published method [86, 94–96]. 8 × 10^6^ cells were distributed equally in 6-well plates in HIV-1 virus-containing media (30mL of 450 ng/mL of HIV-1 p24) and subsequently spinoculated at 480 x *g* for 2 h at 25°C. Cells were then cultured for 5 h in 37°C/5% CO_2_ incubator. After culturing, cells were pelleted by centrifugation for 10 min (300 x *g*). The supernatant was then carefully aspirated, and the cell pellet was washed twice with 2 mL of K−/− buffer (20 mM HEPES [pH 7.6], 150 mM KCl, 5 mM MgCl_2_) at room temperature. The pellet was then gently lysed by resuspending in 2 mL of ice-cold K+/+ buffer (20 mM HEPES [pH 7.6], 150 mM KCl, 5 mM MgCl_2_, 1 mM dithiothreitol [DTT], 20 μg/mL aprotinin, 0.025% [w/v] digitonin) and rocking for 10 min at room temperature. The cytoplasmic extract containing the PICs was then separated from other cellular components by differential centrifugation for 4 min at 1,500 × g at 4°C. The supernatant was transferred to a new microcentrifuge tube and centrifuged for 1 min at 16,000 × g at 4°C. The resulting supernatant was transferred to a new microcentrifuge tube and treated with RNase A (Thermo Fisher Scientific) for 30 min at room temperature to remove any cellular and or viral RNA. Finally, 60% sucrose (w/v) in K−/− buffer was added to a final concentration of 7% and gently mixed by pipetting. These PICs were then aliquoted, flash frozen in liquid nitrogen, and then stored in a −80°C freezer.

### Assay for PIC-associated integration activity *in vitro*

*In vitro* integration assays were performed using a modified version of published protocol [97]. The target DNA used was a PCR-amplified 2-kb region of the phi-X174 genome. The PCR was performed in a final volume of 50 μL containing 50 ng of the phi-X174 plasmid (Promega), 500 nM concentrations (each) of the forward primer (5’-CGCTTCCATGACGCAGAAGTT-3’) and the reverse primer (5’-CACTGACCCTCAGCAATCTTA-3’), 1X Phusion reaction buffer (NEB), dNTP nucleotide mix containing 200 μM concentrations of each nucleotide (NEB), and 1.25 U of Phusion DNA Polymerase (NEB) under the following thermocycling conditions: initial incubation at 95°C for 2 min; followed by 34 cycles at 95°C for 30 s, 53°C for 30 s, and 72°C for 2 min; and a final incubation at 72°C for 10 min. PCR products were resolved by standard agarose gel electrophoresis, and the phiX174-specific PCR amplicon was gel purified by using a Gel DNA Recovery Kit (Zymo Research). The *in vitro* integration reaction was carried out by mixing 200 μL of PICs and 600 ng of target DNA and then incubating the mixture at 37°C for 45 min. The integration reaction was stopped and deproteinized by adding SDS, EDTA, and proteinase K to final concentrations of 0.5%, 8 mM, and 0.5 mg/mL respectively, followed by incubation overnight at 56°C. The deproteinized sample was mixed with an equal volume of phenol (equilibrated with 10 mM Tris HCl), mixed thoroughly by vortexing, and centrifuged (16,000 x *g*) for 2 min at room temperature. The aqueous phase was extracted once with an equal volume of phenol-chloroform (1:1) mixture, followed by an equal volume of chloroform. The DNA was precipitated by adding 2.5 volumes of 100% of ice-cold ethanol in the presence of sodium acetate (0.3 M, final concentration) and the coprecipitant glycogen (25 to 100 μg, final concentration), followed by an incubation overnight at-80°C. The sample was centrifuged (16,000 x *g*) for 30 min at 4°C, and the resultant DNA pellet was washed once with 80% ethanol using centrifugation (16,000 x *g*) for 10 min at 4°C. The precipitated DNA was air dried at room temperature, resuspended in 50 μL of nuclease-free water, and used as the template DNA in a nested qPCR. A first-round standard PCR, designed to amplify only the integrated virus-target DNA junctions, was carried out in a final volume of 50 μL containing 5 μL of purified DNA product from the integration reaction, 500 nM concentrations (each) of primers targeting the target DNA (5’-CACTGACCCTCAGCAATCTTA-3’) and the viral LTR (5’-GTGCGCGCTTCAGCAAG-3’), 2X Megafi Reaction Buffer (ABM), and 1.25 U of MegaFi DNA polymerase (ABM) under the following thermocycling conditions: initial incubation at 95°C for 5 min; followed by 23 cycles at 94°C for 30 s, 55°C for 30 s and 72°C for 2 min; and a final incubation at 72°C for 10 min. The second-round qPCR designed to amplify only the viral LTR-specific region contained a 1/10 volume of the first-round PCR products as the template DNA, 1X iTaq Universal Probe Supermix (Bio-Rad), 300 nM concentrations (each) of the viral LTR-specific primers that target the R region (5’TCTGGCTAACTAGGGAACCCA-3’) and the U5 region (5’-CTGACTAAAAGGGTCTGAGG-3’), and a 100 nM concentration of TaqMan probe (5’-[6-FAM]-TTAAGCCTCAATAAAGCTTGCCTTGAGTGC-[TAMRA]-3’). The qPCR run included an initial incubation at 95°C for 3 min, followed by 39 cycles of amplification and acquisition at 94°C for 15 s and 58°C for 30 s, and 72°C for 30 s. During qPCR of the samples, a standard curve was generated in parallel and under the same conditions using 10-fold serial dilutions of known copy numbers (10^0^ to 10^8^) of the HIV-1 molecular clone plasmid. Data were analyzed using CFX Manager software (Bio-Rad), and integrated viral DNA copy numbers were determined by plotting the qPCR data against the standard curve. To determine the integration efficiency (i.e., ratio of chromosome-integrated viral DNA copy numbers to corresponding PIC-associated viral DNA copy numbers) of the *in vitro* integration reactions, the PIC-associated viral DNA was isolated and copy numbers were determined by qPCR.

### Integration site selection

Genomic DNA (∼5 µg) was digested overnight with a cocktail of enzymes (MseI and BglII; 100 U each). Following overnight ligation (4 parallel reactions) to asymmetric linkers, DNA was purified using a PCR purification kit. The ligated samples were subjected to two rounds of ligation-mediated (LM)-PCR using virus and linker-specific primers. Following PCR purification, libraries were assessed for fragment size distribution by TapeStation-4150, quantified by DNA fluorimetry (Qubit), and then pooled at 10 nM. Pooled samples were further diluted to 2 nM in Illumina sequencing resuspension buffer (RSB, Ref#20762979). Next, PhiX Control v3 DNA was spiked at 30%, and the sample was diluted to 650 pM with RSB. The mixture (20 µl) was loaded into a P1 300 cycle cartridge and sequenced on an Illumina NextSeq 2000 sequencer. Raw fastq files were demultiplexed using Sabre tool or by a Perl script [98]. Post demultiplexing, files were trimmed, aligned to human genome build hg19, and bed files were generated as described[76, 91]. Integration into genes and SPADs was scored as within these genomic coordinates. For TSSs, CpG islands, and LADs, sites were mapped within +/-2.5 kb windows (5 kb surrounding these coordinates). Gene density was assessed as number of genes per Mb. Random integration controls (RICs) were generated by shearing hg19 in silico using the restriction enzyme sites used to generate the wet-bench samples, and then mapping the resultant fragments with respect to aforementioned genomic annotations [91].

### Statistical analysis

All experiments were conducted at least three times with triplicates. Data were expressed as mean ± standard error of mean (SEM) obtained from three independent experiments. Significance of differences between control and treated samples were determined by Student’s *t-*test. The difference between groups were determined by paired, two-tailed student’s t-test. A p-value of < 0.05 was considered statistically significant.

### Data availability statement

All the data generated in this study are included in the manuscript either as main figures or supplementary figures.

## RESULTS

### CPSF6 depletion reduces HIV-1 PIC-associated viral DNA integration activity

CPSF6 binds to the HIV-1 capsid and plays multiple roles during early steps of infection [68]. However, the dominant function of CPSF6 seems to direct the HIV-1 PIC into actively transcribing gene bodies of the host genome for viral DNA integration [82–84]. However, the mechanism by which CPSF6 regulates integration site selection is not fully understood. Particularly, the effects of CPSF6 on PIC function have not been previously reported. PICs are sub-viral nucleoprotein complexes consisting of the viral IN, viral DNA and other viral/host factors that carries out viral DNA integration to establish infection [65]. Importantly, PICs can be extracted from infected cells and these PICs retain viral DNA integration activity *in vitro* [99, 100]. Therefore, to study CPSF6’s effect on PIC function, we first inoculated HEK293T [83] and HeLa CPSF6 knockout (CKO) cells [92], as well as respective control cells, with cell-free DNase I-treated VSV-G pseudotyped HIV-1 particles. Then, cytoplasmic extracts containing the PICs were prepared [86, 94, 97] and subjected to an *in vitro* integration assay containing a linear double-stranded target DNA substrate. Then, the copies of viral DNA integrated into the target DNA were quantified by a nested qPCR method [86, 94, 101]. As expected, the PICs from the infected control cells efficiently carried out viral DNA integration and the integration activity was inhibited by the IN inhibitor Raltegravir (RAL) (Fig. 1A). Moreover, measurable integration activity was not detected in reactions lacking PICs or target DNA, confirming the specificity of the assay. Our results further revealed that PICs from the CKO HEK293T cells exhibited significantly lower integration activity (a four-fold decrease) when compared to the PICs from the control cells (Fig. 1B). Quantification of the viral DNA levels in these PIC preparations revealed lower viral DNA in the PICs from CKO cells compared to the control cells (Fig. 1C). Therefore, we determined PIC specific activity by calculating the ratio of the copies of integrated viral DNA to the copies of PIC-associated viral DNA (Fig. 1D). These calculations indicated that the specific integration activity of PICs from CKO HEK293T cells was significantly lower (∼40%) compared to those from the control cells. Similar defects in PIC activity were exhibited using extracts of CKO HeLa cells (Fig. 1E-G). For example, PICs from CKO HeLa cells showed a significantly lower integration activity (almost ∼4-5 fold decrease) when compared to the PICs from the control cells (Fig. 1E). The viral DNA levels in the PICs of the WT and CKO HeLa cells were minimally changed (Fig. 1F), in contrast to the lower viral DNA levels in the HEK293T CKO cells (Fig. 1C). Nevertheless, PIC specific activity calculations revealed that integration activity of PICs from the CKO HeLa cells was significantly lower when compared to the PICs from the control cells (Fig. 1G). Collectively, these biochemical studies demonstrate that PICs extracted from CPSF6-depleted cells exhibit lower viral DNA integration activity in vitro, thus suggesting that CPSF6 is critical for HIV-1 PIC function.

**Figure 1.**
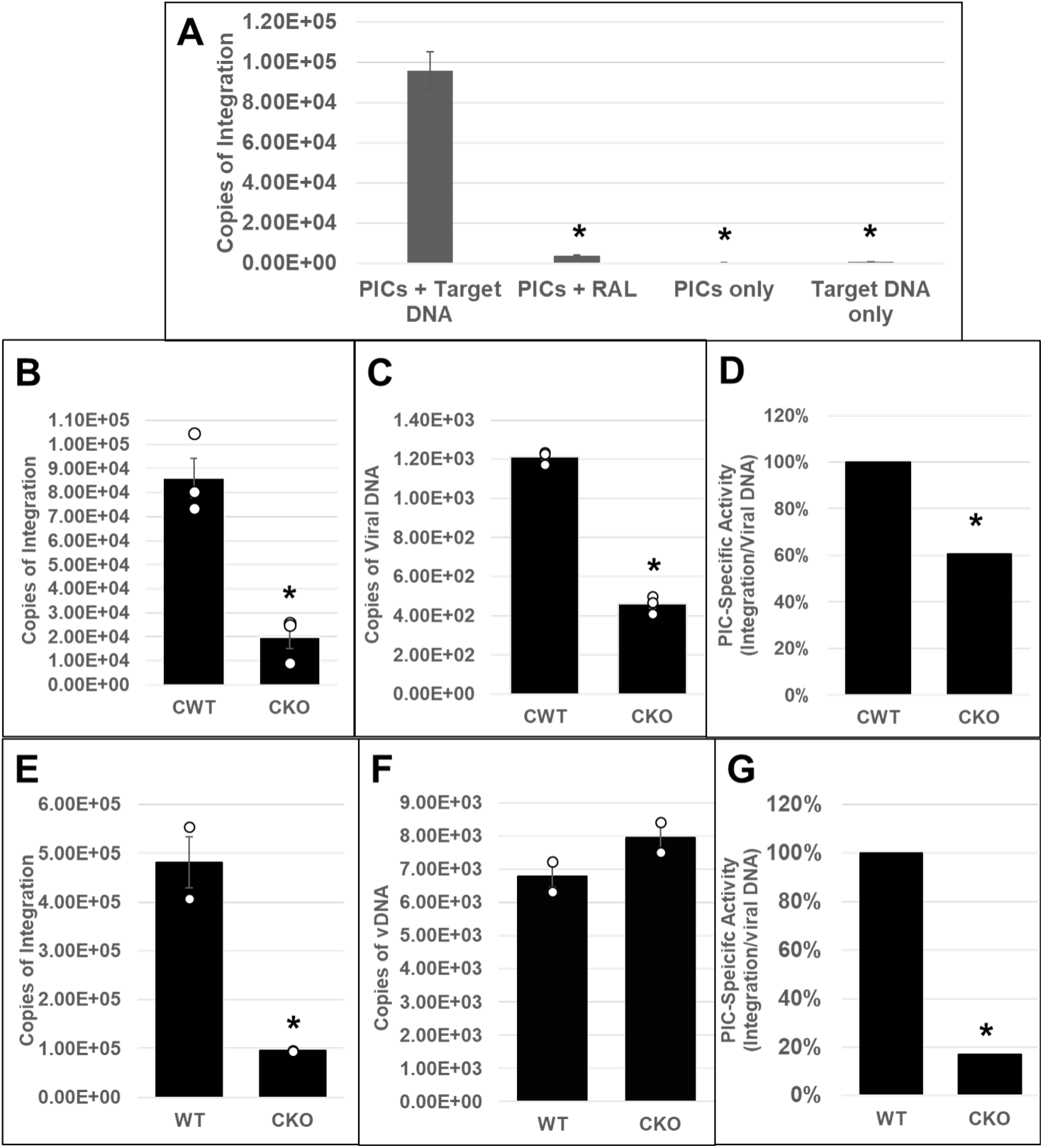
Effect of CPSF6-depletion on HIV-1 PIC integration activity in vitro. (A-D) HEK293T CWT and CKO cells were spinoculated with 450 ng p24 virus/million cells as described in methods. PICs were extracted and in vitro integration reactions were carried out using phiX174 target DNA. Reactions in the presence of the IN inhibitor raltegravir (RAL) or in the absence of the target DNA served as controls. A nested qPCR-based approach was utilized to measure the integration activity of the PICs. **(A)** Assessment and verification of PIC integration activity. **(B)** Comparison of in vitro integration activity of PICs extracted from CWT or CKO cells. **(C)** Copies of PIC-associated viral DNA determined via qPCR. **(D)** PIC specific activity normalized to the amount of viral DNA in the different preps (integration/viral DNA). Specific activity is displayed as percentage relative to CWT cell PICs. **(E-G)** PIC activity from CKO and WT HeLa cells. **(E)** PIC-associated integration activity. **(F)** Viral DNA content of the PICs. **(G)** Specific PIC activity. White circles in graphs represent separate biological replicates. Error bars represent SEM. The p-values (*) represents statistical significance (p< 0.05) between control and CKO PICs.

### Disruption of CPSF6-CA interaction reduces HIV-1 PIC-associated viral DNA integration activity

Our studies of PICs from CPSF6-depleted HEK293T and HeLa cells supported a functional role of CPSF6 in PIC activity. However, these cell types are not permissive to HIV-1 infection. Therefore, we next sought to study PICs using a T cell line model, which is arguably more physiologically relevant for HIV-1 studies. First, we employed a CRISPR-Cas9 based approach to deplete CPSF6 in Jurkat cells, a T cell line that is both permissive to HIV-1 infection [53] and has been used for PIC extraction in our laboratory [101, 102]. Unfortunately, we were unable to generate CKO Jurkat cells and only partial depletion of CPSF6 was achieved after several attempts (data not shown). Efforts to isolate individual clones of CKO Jurkat cells from these polyclonal cell pools by serial dilution were also unsuccessful. Previous study also reported challenges associated with generating CKO Jurkat cells [91, 103]. Given this, we next applied a CRISPR-Cas9-based microhomology-mediated end-joining strategy [104] to mutate the region of *CPSF6* exon 7 encoding the CA-binding phenylalanine-glycine (FG) motif to alanine-alanine (AA) (Suppl. Fig. 1A-B). Following nucleofection of crRNPs and a single-stranded DNA repair template, cells were cloned by limiting dilution and individual cell clones were expanded and analyzed for the site-directed mutations. A second set of cell clones was derived under identical conditions using non-targeting (NT) guide RNA as controls. Characterization of the SupT1 WT and CPSF6 knock-in (CKI) cell lines showed minimal differences in CPSF6 expression (Suppl. Fig. 1C) and cell viability (Suppl. Fig. 2). This recoding approach has also been used to generate CKI Jurkat and CEM T cells, the results of which will be presented elsewhere. However, due to the long-standing usage of SupT1 cells in HIV-1 PIC studies [66, 67, 86, 94–96, 99, 100, 105, 106], the SupT1 WT and CKI clonal cell lines were selected for this study.

PICs extracted from WT and CKI clonal cell lines inoculated with HIV-1 particles were subjected to in vitro integration assays. Measurement of viral DNA integration revealed that the PICs from the WT SupT1 cells efficiently integrated the viral DNA into the exogenous target DNA and this activity was significantly reduced in the presence of RAL (Fig. 2A). Notably, the PICs from the WT SupT1 cells exhibited significantly higher integration activity-almost 2-log-fold higher, when compared to the PICs from CPSF6 WT (CWT) HEK293T cells (Fig. 1A) and CWT HeLa cells (Fig. 1E). However, the PICs from the CKI SupT1 cells exhibited significantly lower integration activity (∼2-log defect) when compared to the PICs from the WT SupT1 cells (Fig. 2B). This level of reduction was in stark contrast to the ∼5-fold reductions in integration activity observed with the PICs from CKO HEK293T and CKO HeLa cells (Fig. 1B and E). To assess whether the lower integration activity of PICs from CKI SupT1 cells was due to a reduction in the number of PICs formed in the CKI cells, we quantified the amount of viral DNA in the PIC preparations. We found that the viral DNA levels in the CKI cells were not significantly different from that of the control cells (Fig. 2C). Accordingly, estimation of PIC specific activity by calculating the ratio of the copies of integrated viral DNA to the copies of PIC-associated viral DNA indicated that the integration activity of PICs of CKI cells was significantly lower (>95%) compared to the control cell complexes (Fig. 2D). Together, our results in Figures 1 and 2 provide in vitro biochemical evidence that PICs from cells depleted of CPSF6 or cells that express a mutant CPSF6 that cannot bind CA exhibit significantly lower integration activity.

**Figure 2.**
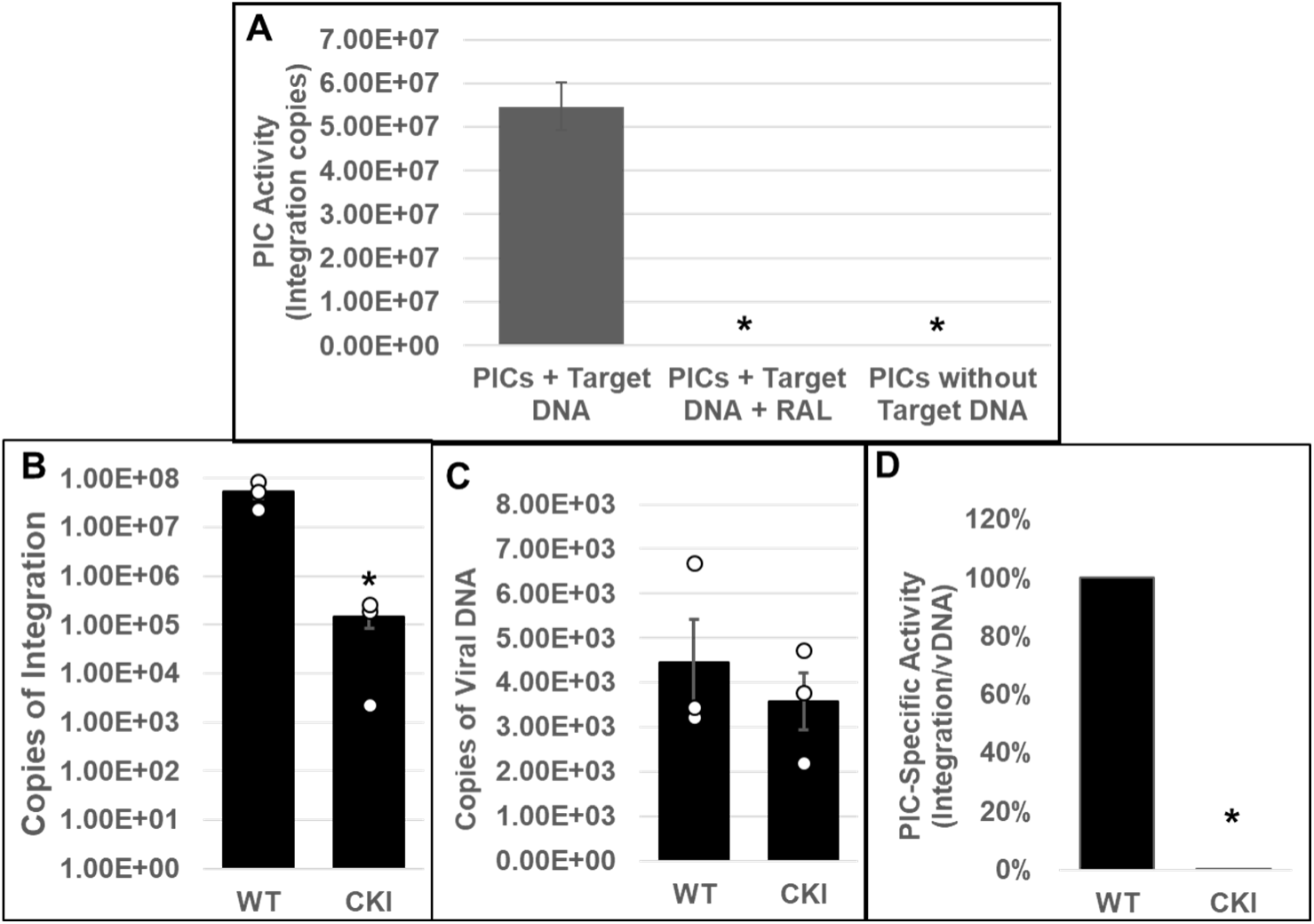
Effect of disrupting the CPSF6-CA interaction on the in vitro integration activity of HIV-1 PICs. **(A)** Parental WT SupT1 cells or CKI cells were spinoculated with high titer infectious HIV-1 particles (1500 ng p24/mL) and cultured for 5 h followed by extraction of PIC-containing cytoplasmic extracts. *In vitro* assays using these PICs as the source of integration activity and quantification of viral DNA integration into an exogenous target DNA by nested PCR were carried out with appropriate controls. **(B)** In vitro integration assays of PICs extracted from WT1 and from CKI cells were carried out and the copy numbers of integrated viral DNAs were determined. **(C)** Viral DNA copy numbers were quantified by qPCR. **(D)** The specific activity of the PIC-mediated integration was determined by calculating the ratio of integrated viral DNA copy numbers (from panel B) to the corresponding PIC-associated viral DNA copy numbers (from panel C). Mean values from two independent experiments, each conducted in triplicate are shown with error bars representing the SEM. White circles present in graphs represent individual data points. The p-values (*) represents statistical significance (p< 0.05) between the control and CKI PICs.

### Addition of CPSF6 enhances the integration activity of HIV-1 PICs

Our integration activity measurements suggested that CPSF6 is important for PIC function (Figs. 1-2). To pinpoint whether CPSF6 directly regulates PIC-mediated viral DNA integration, we examined the effect of the addition of purified recombinant CPSF6 protein to in vitro integration reactions (Fig. 3). Purified recombinant CPSF6 protein was added at increasing concentrations (0, 0.5, 1, and 2 μM) to the reaction mixtures containing PICs extracted from WT or CKI SupT1 cells. Quantification of viral DNA integration showed that addition of CPSF6 significantly stimulated the activity of PICs from both the WT (Fig. 3A-B) and CKI cells (Fig. 3C-D). Because of larger number of PIC samples to be analyzed, we used 1/10^th^ volume of PICs in this assay compared to the previous assays to reduce batch to batch variation in PIC activity. Therefore, the integration copies in this assay were expectedly lower than those measured in Fig. 2A. Nevertheless, the WT PICs showed markedly higher activity-a log-fold higher, when compared to the CKI PICs, which is consistent with the results in Figures 1 and 2. Addition of 0.5 μM CPSF6 enhanced the integration activity of WT PICs by ∼2.5-fold, whereas 1.0 μM CPSF6 increased it by almost 4-fold (Fig. 3B). However, the addition of 2.0 μM CPSF6 led only to a minimal increase in the integration activity of WT PICs, suggesting a saturating effect of the added CPSF6 between concentrations of 1 to 2 μM. In contrast, CKI cell-derived PICs showed a dose-dependent increase in integration activity up to the addition of 2 μM CPSF6 (Fig. 3C). Addition of 0.5 μM CPSF6 to the CKI PICs stimulated integration activity by ∼2-fold, a level comparable to that of the control PICs. Moreover, a 3-fold stimulation in integration activity of CKI PICs was observed with 1.0 μM CPSF6 (Fig. 3D), which is slightly lower than that observed with the control PICs. By contrast to the WT condition, addition of 2.0 μM CPSF6 resulted in a 7-fold higher integration activity of CKI PICs. Collectively, these biochemical studies provide additional evidence for a direct and stimulatory role of CPSF6 in PIC function.

**Figure 3:**
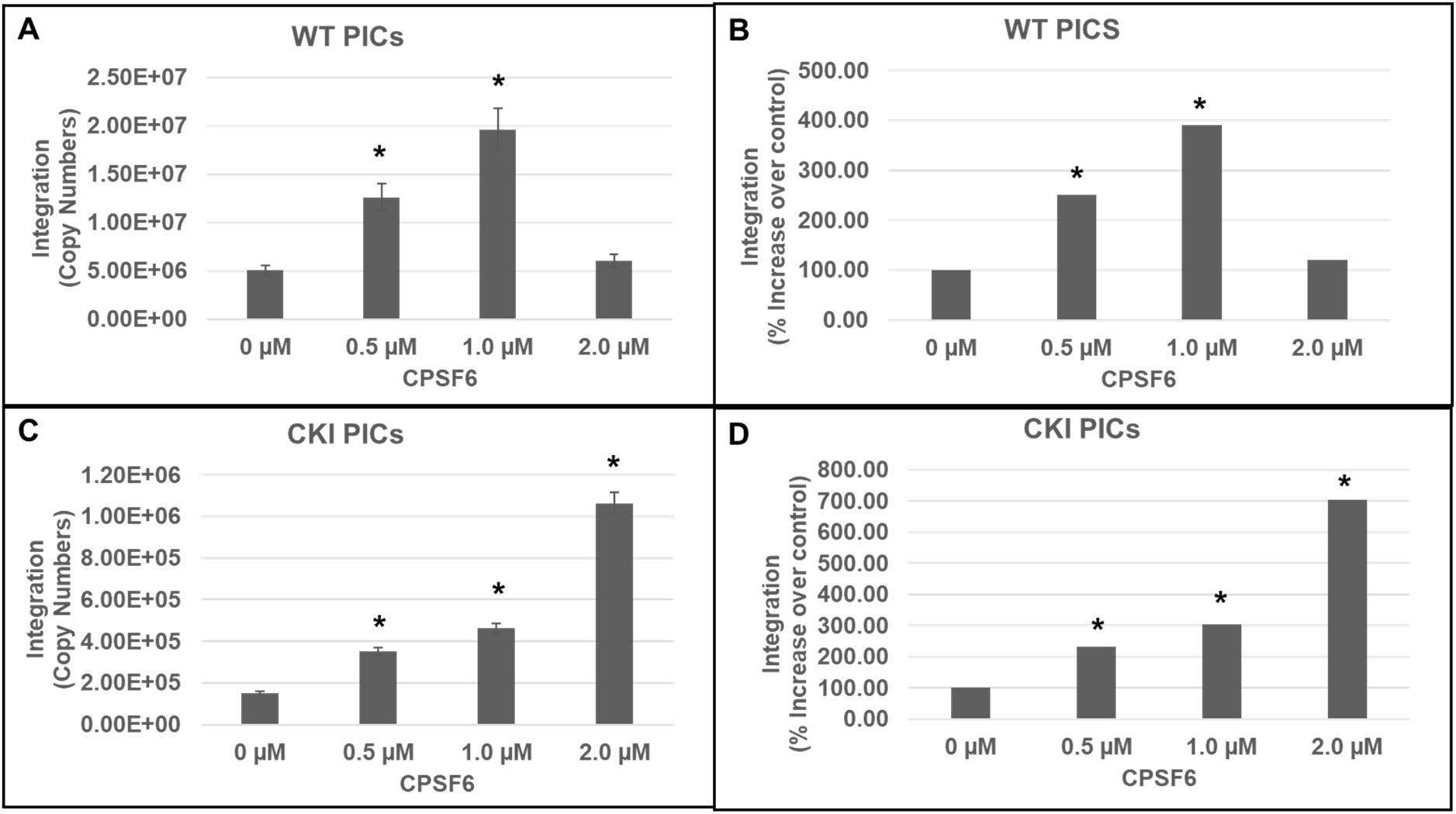
Effects of CPSF6 protein on PIC-associated integration activity. In vitro integration activity of PICs extracted from WT control cells **(A-B)** and from CKI cells **(C-D)** were assessed in the absence or presence of varying concentrations (0-2 µM) of purified recombinant CPSF6 protein. **(A and C)** The copy number of the integrated viral DNAs were determined by nested qPCR. **(B and D)** Integration activity plotted as fold change relative to the integration activity of the respective control PICs. Mean values from two independent experiments, each conducted with triplicate samples, are shown with error bars representing the SEM. The p-values (*) represents statistical significance (p<0.05) relative to untreated PICs (0 µM).

### Disruption of CPSF6-CA interaction blocks HIV-1 DNA integration into the host genome

Our biochemical studies provided in vitro evidence for a critical role of CPSF6 in PIC-mediated viral DNA integration. The PIC is the subviral biochemical machinery that carries out viral DNA integration into the host genome [65]. Therefore, to study the effect of CPSF6 on PIC function in an infected cell, we chronicled viral DNA synthesis and integration in the infected WT and CKI clonal cell lines. The WT and CKI SupT1 cells were individually inoculated with pseudotyped HIV-1 particles, and the total DNA isolated from these cells were used to measure various forms of HIV-1 DNA, including integrated DNA detected via Alu-based nested qPCR [97, 107, 108]. Results from these analyses showed a significant reduction (∼75%) in the copies of integrated viral DNA in the CKI cells compared to the WT cells (Fig. 4A-B). These results suggest that CPSF6-CA interaction promotes PIC-mediated viral DNA integration in the context of HIV-1 infection.

**Figure 4:**
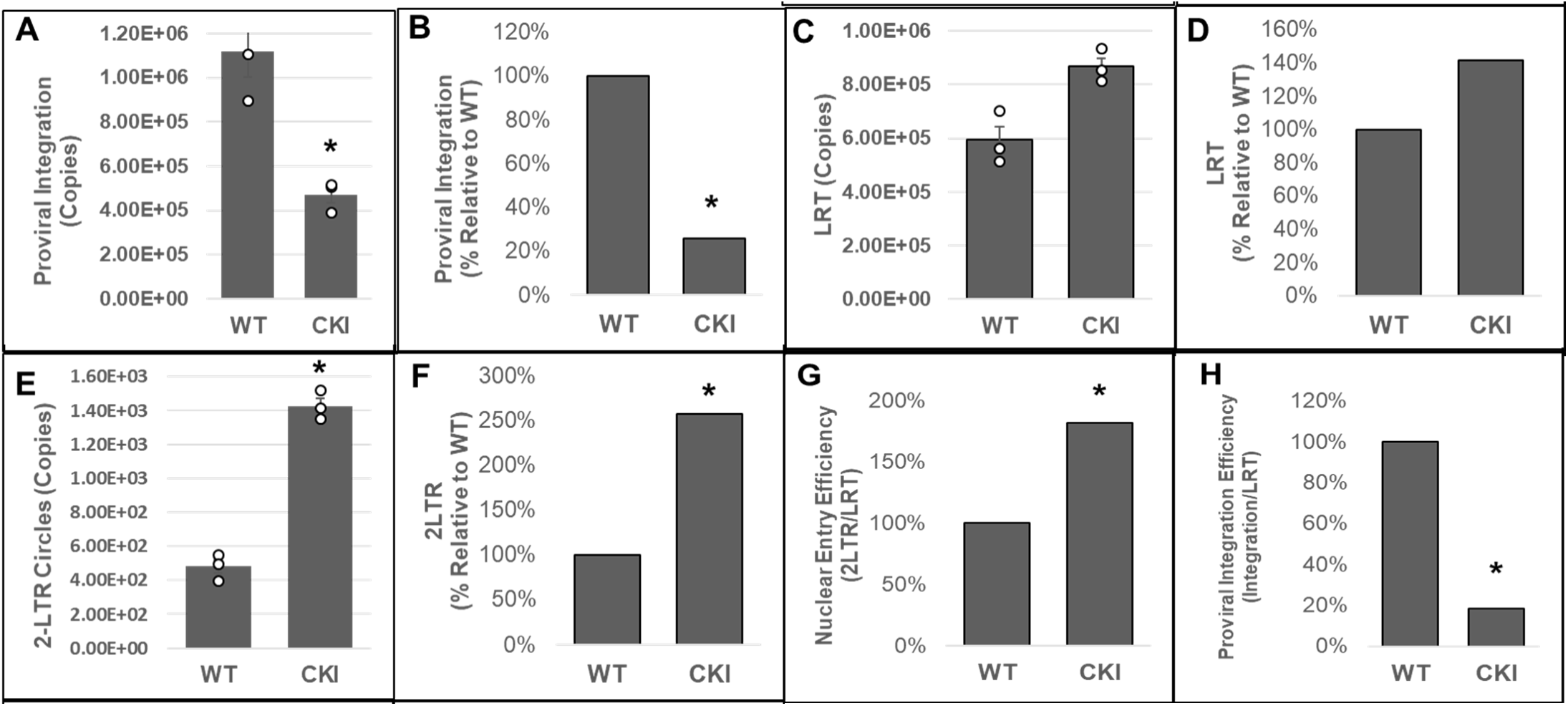
Disruption of the CPSF6-CA interaction blocks HIV-1 integration. **(A-B)** Integrated viral DNA was quantified by *Alu-gag* nested PCR using the total DNA extracted from infected CKI and WT control cells. Copy numbers were calculated from standard curves generated in parallel of known copy numbers (10^0^ - 10^8^) of a HIV-1 molecular clone during the 2^nd^ round qPCR. **(A)** Copies of proviral integrants and **(B)** Percentage of integrants relative to WT1 control cells**. (C-D)** Copies of late reverse transcription (LRT) products were measured by qPCR using a standard curve generated with known copy numbers of a HIV-1 molecular clone. **(C)** Copies of LRT and **(D)** Percentage of LRT relative to the WT1 control cells**. (E-F)** 2-LTR circles were quantified from standard curves generated in parallel using known copy numbers of the p2LTR plasmid. **(E)** Copies of 2-LTR circles and **(F)** Percentage of 2-LTR circles relative to the WT1 control cells**. (G)** A ratio of 2-LTR circles to LRT copies was calculated and plotted as relative percentage to WT control cells. **(H)** A ratio of integration copies-to-LRT was calculated and plotted as relative percentage to the WT control cells. Data are mean values from three independent experiments, each conducted in triplicate with error bars representing the SEM. The p-values (*) represents statistical significance (p< 0.05) between the control and CKI cells. White circles present in graphs represent individual data points.

HIV-1 integration is dependent on the completion of the prior steps of reverse transcription and nuclear import [65]. Notably, HIV-1 CA is known to regulate both reverse transcription and post-reverse transcription steps of infection [16, 52, 55, 84] and the CPSF6-CA interaction is implicated in nuclear entry and post-nuclear entry steps of infection [76, 79, 82, 84, 109–111]. Therefore, we investigated whether the integration defect in the CKI cells was a consequence of a block at either the reverse transcription or nuclear entry step. Copies of late reverse transcription (LRT) products were measured by qPCR using the total DNA from the cells. These measurements revealed that viral DNA synthesis was not reduced in the CKI cells when compared to the WT cells, rather was slightly elevated (∼40%) (Fig. 4C-D). These results suggested that the disruption of CPSF6-CA interaction does not reduce reverse transcription, and the integration block is likely manifested at a post-reverse transcription step. Therefore, we next quantified the levels of 2-LTR circles, a commonly used surrogate marker of nuclear import of retroviral DNA that are generated in the nucleus by non-homologous end-joining [112–116]. Copy numbers of 2-LTR circles in infected CKI cells as well as the WT control cells were calculated by extrapolating qPCR data to a standard curve generated using a plasmid containing the HIV-1 2-LTR junction sequence. Unexpectedly, we observed an increase (∼2.5-fold) in the number of 2-LTR circles in the CKI clones relative to the WT cells (Fig. 4E-F). Moreover, calculation of the efficiency of nuclear entry (ratio of late RT copies to 2-LTR copies) revealed that the nuclear entry of HIV-1 DNA was not reduced in CKI cells, but was seemingly stimulated (Fig. 4G), an observation similar to the reverse transcription in these cells. Finally, calculation of the efficiency of integration (ratio of integration copies to late RT copies) revealed a reduction (∼80%) in integration efficiency in the CKI cells (Fig. 4H). These results suggest that CPSF6 plays an important role in PIC-mediated viral DNA integration in infected cells.

### Disruption of CPSF6-CA interaction minimally affects integration of HIV-1 CA mutant viruses deficient in CPSF6 binding

To solidify the data implicating CPSF6 in HIV-1 integration, we probed the early steps of infection of HIV-1 CA mutant viruses N74D and A77V, which are deficient for CPSF6 binding [77, 78]. We inoculated CKI and control cells with pseudotyped luciferase reporter N74D (Fig. 5A) or A77V (Fig. 5G) viral particles. Twenty-four hours thereafter, luciferase activity was measured in the cellular lysates as an indicator of infectivity. As expected, the infectivity levels of the mutant viruses in CKI cells were comparable to those in control cells, confirming that the infectivity of these mutants is not dependent on CPSF6 binding. Then, to assess the early steps of virus infection, we measured the levels of LRT, 2-LTR circles and viral DNA integration in these cells by qPCR. These measurements indicated that the copies of LRT synthesized by N74D (Fig. 5B) or A77V (Fig. 5H) were not reduced in CKI cells when compared to the control cells. Moreover, there was a slight increase in the LRT levels of these CA mutants in the CKI cells, an effect similar to that observed for WT HIV-1 in CKI cells (Fig. 4C-D). We also observed that the number of 2-LTR circles formed by the mutant viruses in the infected CKI cells was comparable to those formed in the control cells (Fig. 5C and Fig. 5I). Calculation of the nuclear import efficiency indicated a slight but non-significant decrease in nuclear import of the N74D virus in the CKI cells when compared to the control cells (Fig. 5D). However, no such reduction in nuclear import efficiency was detected with the A77V virus in the CKI cells (Fig. 5J). Finally, estimation of the copies of viral DNA integration revealed that integration of N74D (Fig. 5E) or A77V (Fig. 5K) virus was not reduced in the CKI cells when compared to the control cells. Estimation of integration efficiency indicated a slight decrease in N74D integration in CKI cells (Fig. 5F) but no change in the case of the A77V mutant virus (Fig. 5L). The level of decrease in N74D integration (∼20%) matched the reduction in nuclear import (∼20%) in the CKI cells, suggesting that the integration defect was most likely due to the reduction in nuclear import. Together, these results suggest that HIV-1 CA mutants incapable of binding CPSF6 are not blocked for viral DNA integration step in CKI cells, thus providing additional evidence supporting an important role for CPSF6 in HIV-1 integration.

**Figure 5.**
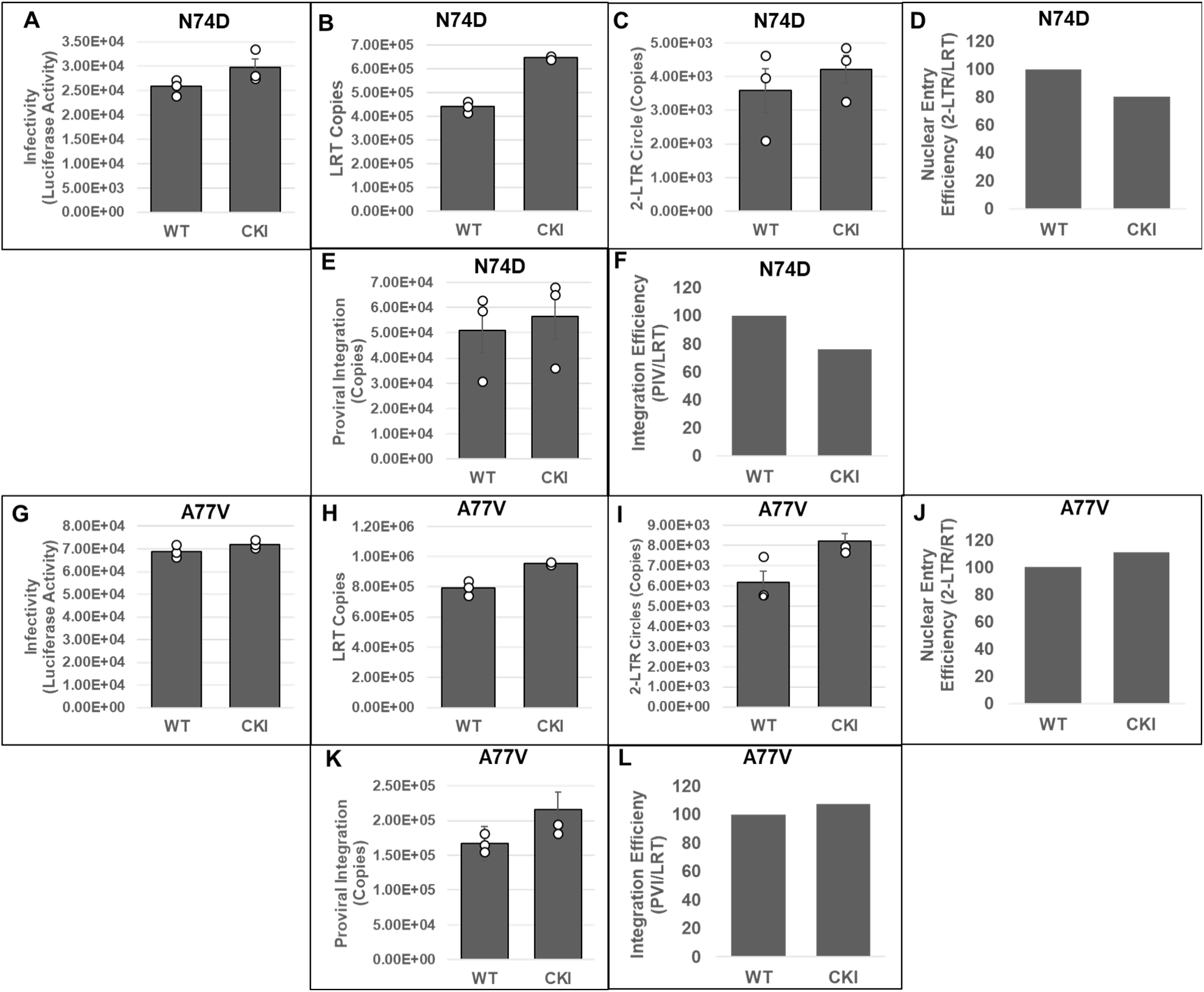
Measurement of early infection steps of HIV-1 CA mutant viruses disrupted for CPSF6 binding. (A and. **G)** Infectivity measurements. WT and CKI clones inoculated with WT CA or mutant CA HIV-1 particles (N74D-panels A-F and A77V-panels G-L) were cultured for 24 h and luciferase activity was measured in the cellular lysates. **(B and H)** Quantification of LRT copies as a measure of reverse transcription. **(C and I)** Quantification of 2-LTR circle copies as a measure of nuclear entry. **(D and J)** Normalized levels of nuclear entry. **(E and K)** Integrated viral DNA copies were quantified by *Alu-gag* nested PCR assay as described in Fig. 1. **(F and L)** A ratio of the copies of proviral integration to LRT copies as a measure of integration efficiency. Data are plotted as relative percentage to the WT control cells. Data are mean values from at least three independent experiments, each conducted in triplicates with error bars representing the SEM. White circles present in graphs represent individual data points.

### Disruption of the CPSF6-CA interaction retargets HIV-1 DNA integration away from SPADs and into LADs

The CPSF6-CA interaction is important for viral cores to move distal from the nuclear rim, the consequences of which help to target viral DNA integration into gene-rich SPADs [76, 79, 83, 84, 91, 117]. Accordingly, CPSF6 depletion or disruption of CPSF6-CA binding misdirects PIC-mediated integration into LADs, which are proximal to the nuclear envelope. Our results indicated that CPSF6 is critical for HIV-1 DNA integration, both in vitro (Fig. 1-3) and in infected cells (Fig. 4). Therefore, to further understand the role of CPSF6 in PIC function, we mapped proviral integration sites in CKI and control cells. Genomic DNA from infected cells were sheared and ligated to asymmetric linkers and the resulting DNAs were purified. Then, LTR-linker sequences were amplified via a semi-nested PCR assay and were analyzed by Illumina sequencing [118, 119]. Unique sites of HIV-1 integration were then mapped with respect to several genomic features, including genes, SPADs, LADs, CpG islands, transcription start sites (TSS) and gene dense regions, as well as computationally-generated random integration control (RIC) values (Fig. 6). In control cells, as expected, HIV-1 integration into genes was significantly enriched when compared to the RIC value (Fig. 6A). However, in the CKI cells, there was a significant reduction in integration into genes (Fig. 6A). Notably, the integration sites within the CKI cells were significantly retargeted away from SPADs, relative to the integration sites in the control cells (Fig. 6B). For example, SPAD-proximal integration in WT cells was approximately 35%, whereas in the CKI cells it was significantly reduced to ∼0.5% (Fig 6B), a value significantly lower than the 4.8% RIC value for SPAD-proximal integration. Expectedly, there was a concurrent and significant increase in integration into LADs in the CKI cells (∼55%), whereas, not surprisingly, integration into LADs was significantly lower in the WT1 cells (Fig. 6C). Furthermore, while 1.7% and 2.3% integrations were within 2.5 kb of CpG island and TSS, respectively, in CKI cells, 6% and 4.6% integrations were within the CpG island and TSS, respectively, in control cells (Fig. 6D-E). Finally, integrations into gene-dense regions of chromatin were significantly reduced in CKI cells, from about 21 genes/Mb in WT cells to 7 genes/Mb, which was significantly lower than the 8.9 genes/Mb RIC value (Fig. 6F). These integration site analyses indicate that disrupting CPSF6-CA interaction profoundly redirects the PIC away from SPADs towards the low-gene dense regions of the LADs near the nuclear envelope for viral DNA integration.

**Figure 6.**
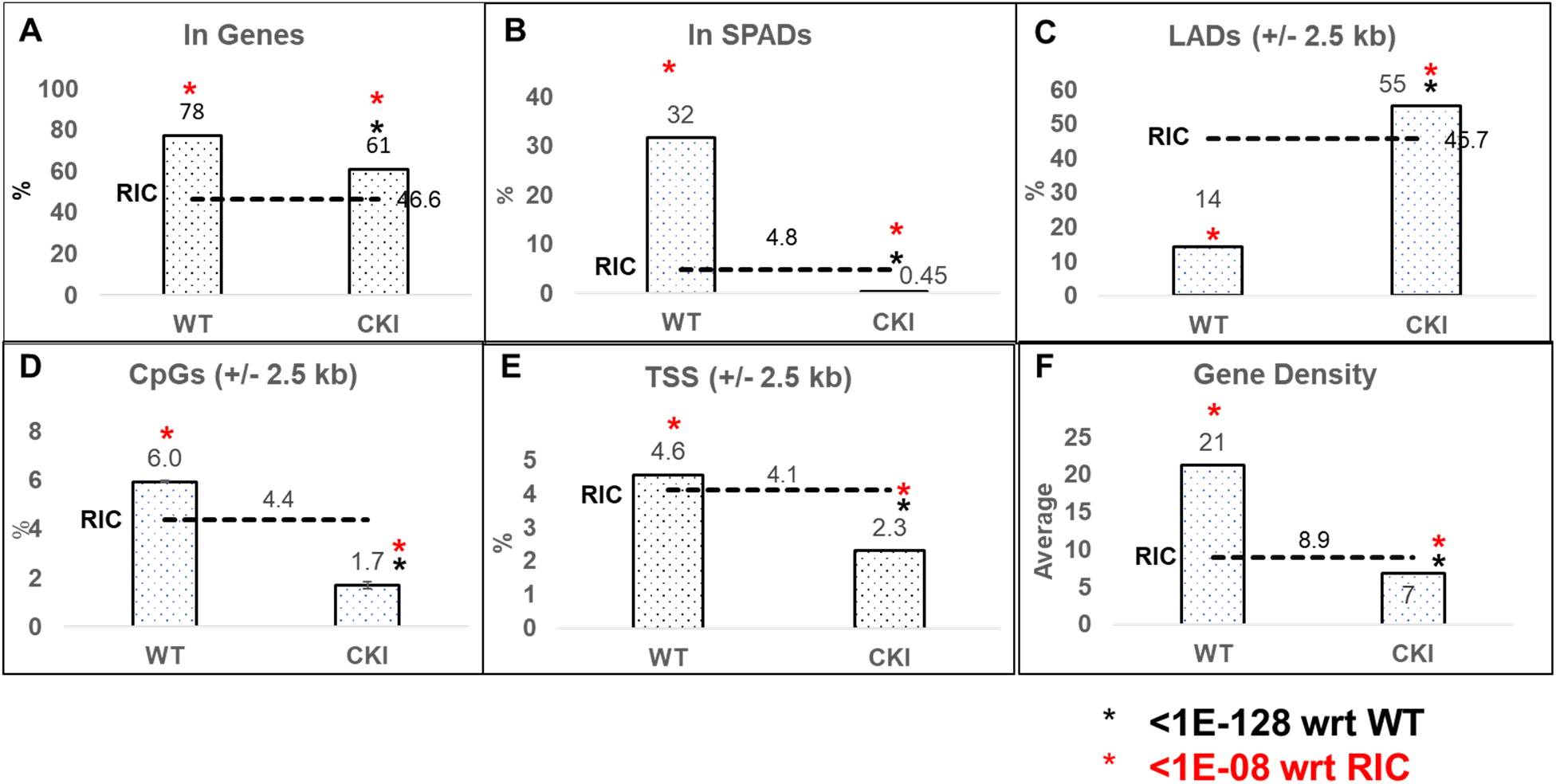
Disruption of CPSF6-CA Interaction retargets HIV-1 DNA away from SPADs and into LADs. Integration into **(A)** genes and **(B)** SPADs was scored as within these genomic coordinates. For **(C)** LADs, **(D)** CpG islands, and **(E)** TSSs, sites were mapped within +/-2.5 kb windows (5 kb surrounding these coordinates). **(F)** Gene density within the 500 kb surrounding integration sites. Gene density was assessed as number of genes per Mb. Random integration controls (RICs) were generated by shearing hg19 in silico using the restriction enzyme sites used to generate the wet-bench samples, and then mapping the resultant fragments with respect to aforementioned genomic annotations. The p-values (*) represent statistical significance with respect to (wrt) WT cells (black asterisks; <10^-128^) and matched RIC values (red asterisks; <10^-8^).

### Effect of CPSF6-CA interaction on HIV-1 infectivity is context dependent

Our in vitro and in cell studies of HIV-1 integration strongly suggested that CPSF6 promotes HIV-1 PIC function. To probe how the effects of CPSF6 on PIC function influenced HIV-1 infectivity, we carried out single-round infection studies in both CKO and CKI cells. First, we inoculated CKO and CWT HEK293T cells with three different amounts of pseudotyped luciferase reporter HIV-1 particles. 24 or 48 h post-infection (hpi), luciferase activity was measured in the cellular lysates as an indicator of virus infectivity. Results from these assays showed that, compared to the CWT cells, HIV-1 infection was stimulated in CKO cells at both 24 hpi (Fig. 7) and 48 hpi (Suppl. Fig. 3). The increased infectivity in the CKO cells was sustained at all the three concentrations of virus used for inoculation (Fig. 7 and Suppl. Fig. 3). These infectivity results are consistent with previously reported data from studies using these cells [83]

**Fig. 7.**
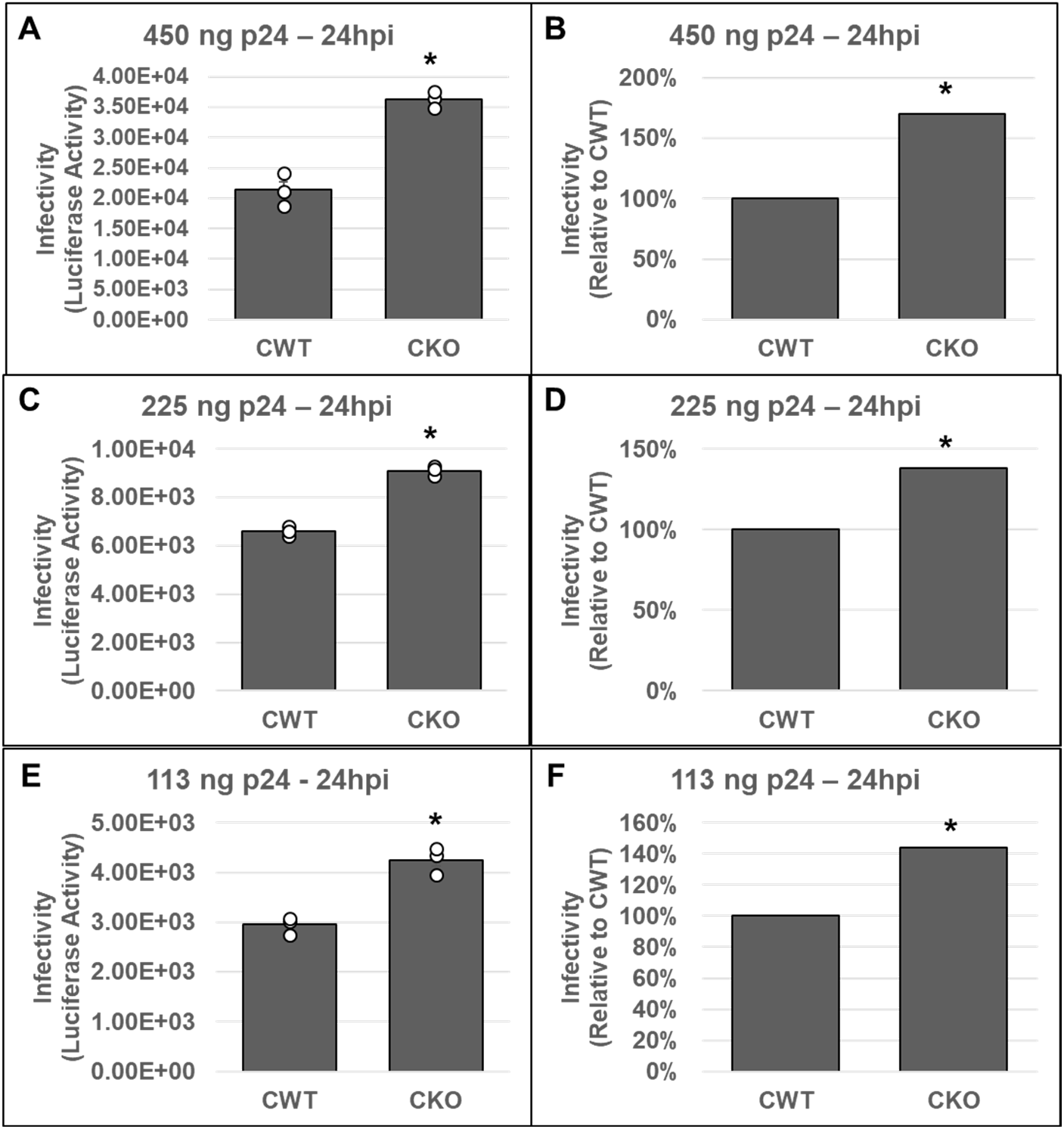
Effects of CPSF6 depletion on HIV-1 infectivity. CPSF6 knockout (CKO) and control HEK293T (CWT) cell lines were inoculated with three different concentrations of pseudotyped HIV-1.Luc virus as described in methods; the total infection timecourse was 24 h. Cell pellets were then collected and subjected to infectivity evaluation by measurement of luciferase activity. Infectivity is displayed as both luciferase activity (**A, C, E)** and as a percentage relative to the luciferase activity of the corresponding CWT control **(B, D, F)**. Three virus concentrations included **(A, B)** 450 ng p24, **(C, D)** 225 ng p24, and **(E, F)** 113 ng p24. Data are representative of three independent experiments each conducted in triplicate with error bars representing the SEM. The p-values (*) represents statistical significance (p< 0.05) CWT and CKO cells. White circles present in graphs represent individual data points.

Next, three parental (WT control) and two CKI SupT1 clones were individually inoculated with pseudotyped luciferase reporter HIV-1 particles and virus infectivity was measured at 24 or 48 hpi. These measurements showed that the WT control cells supported higher virus infectivity when compared to the CKI clones at 24 hpi (Suppl. Fig. 4A). Normalization of the luciferase activity to the total protein levels or to the viability of the respective infected cells minimally changed the level of reduction in virus infectivity in the CKI clones (Suppl. Fig. 4B-C). To comprehensively probe the virus infectivity levels in the CKI clones, we performed pairwise matched infections of WT versus CKI cell lines (Fig. 8). Results from these analyses showed reduction in viral infectivity in both CKI cell lines at 24 hpi, when compared across control cell lines. Particularly, when compared to WT1 control cells, an estimated 70% and 60% decrease in infectivity was observed in CKI7 (Fig. 8A-B) and in CKI19 (Fig. 8C-D) cells, respectively. However, when compared to WT2 cells, an estimated 30% decrease in infectivity was observed for both CKI7 (Fig. 8E-F) and CKI19 (Fig. 8G-H) cells. Conversely, when compared to WT3 cells, an estimated 80% and 70% decrease in infectivity was observed for CKI7 (Fig. 8I-J) and CKI19 (Fig. 8K-L) cells, respectively. Considering the large clonal-based variation in infectivity levels in these recoded SupT1 cells, we compared the aggregate infectivity data of all WT and CKI cell lines (Fig. 8M-N). These results revealed that HIV-1 infectivity was significantly reduced by ∼50% across CKI cells compared to the WT cells. Notably, by 48 hpi, the CKI infection defects noted at 24 hpi had dissipated, both at the level of cell type paired match infections (Fig. 9A-L) and aggregate cell infection data (Fig. 9M-N). A minimal change in HIV-1 infectivity at 48 hpi was previously reported in CPSF6-depleted Jurkat cells in single-round infection assays [91].

**Figure 8:**
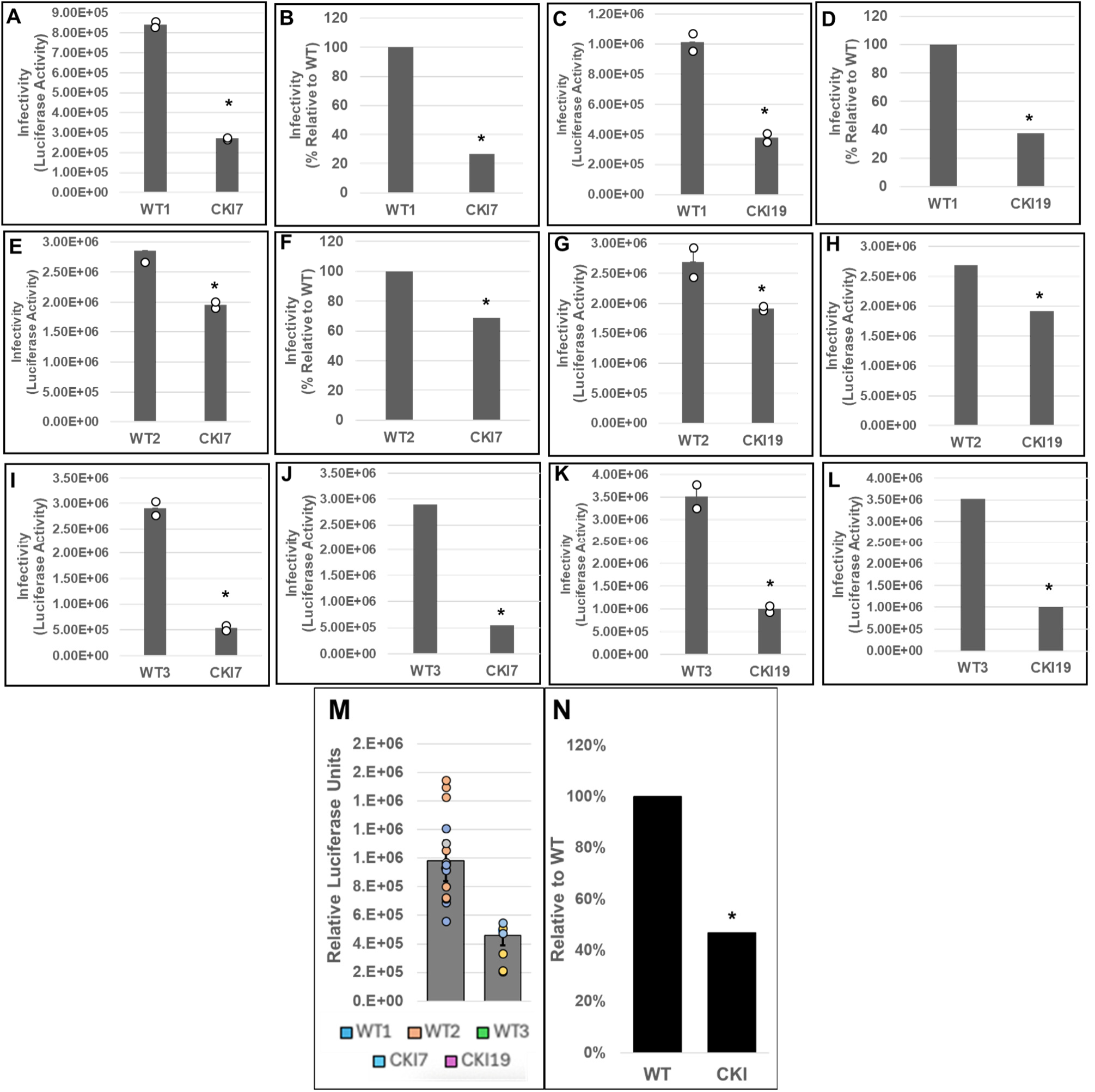
Effects of disrupting the CPSF6-CA interaction on HIV-1 infection at 24 hpi. WT cell lines WT1 **(A-D)**, WT2 **(E-H)** and WT3 **(I-L)** and CKI SupT1 cell lines CKI7 **(A-B, E-F, and I-J)**, and CKI19 **(C-D, G-H, and K-L)** were spinoculated with 450 ng p24 of pseudotyped HIV-1.Luc reporter particles. The total infection timecourse was 24 h, and luciferase activity was measured in the cellular lysates as an indicator of infectivity (Panels-A, C, E, G, I, and K). Infectivity data were also plotted as percent infectivity relative to respective control clones (Panels-B, D, F, H, J, and L). Data shown are mean values from three independent experiments, each conducted in triplicates. **(M)** Infectivity values sorted by editing treatment (CPSF6 non-edited vs edited). Colored circles individual data point from corresponding cell line. **(N)** Displays aggregate infectivity as a percent relative to WT. Error bars represent the standard error of the mean (SEM) and the p-values (*) represents statistical significance (p< 0.05) between the control and CKI cells.

**Figure 9:**
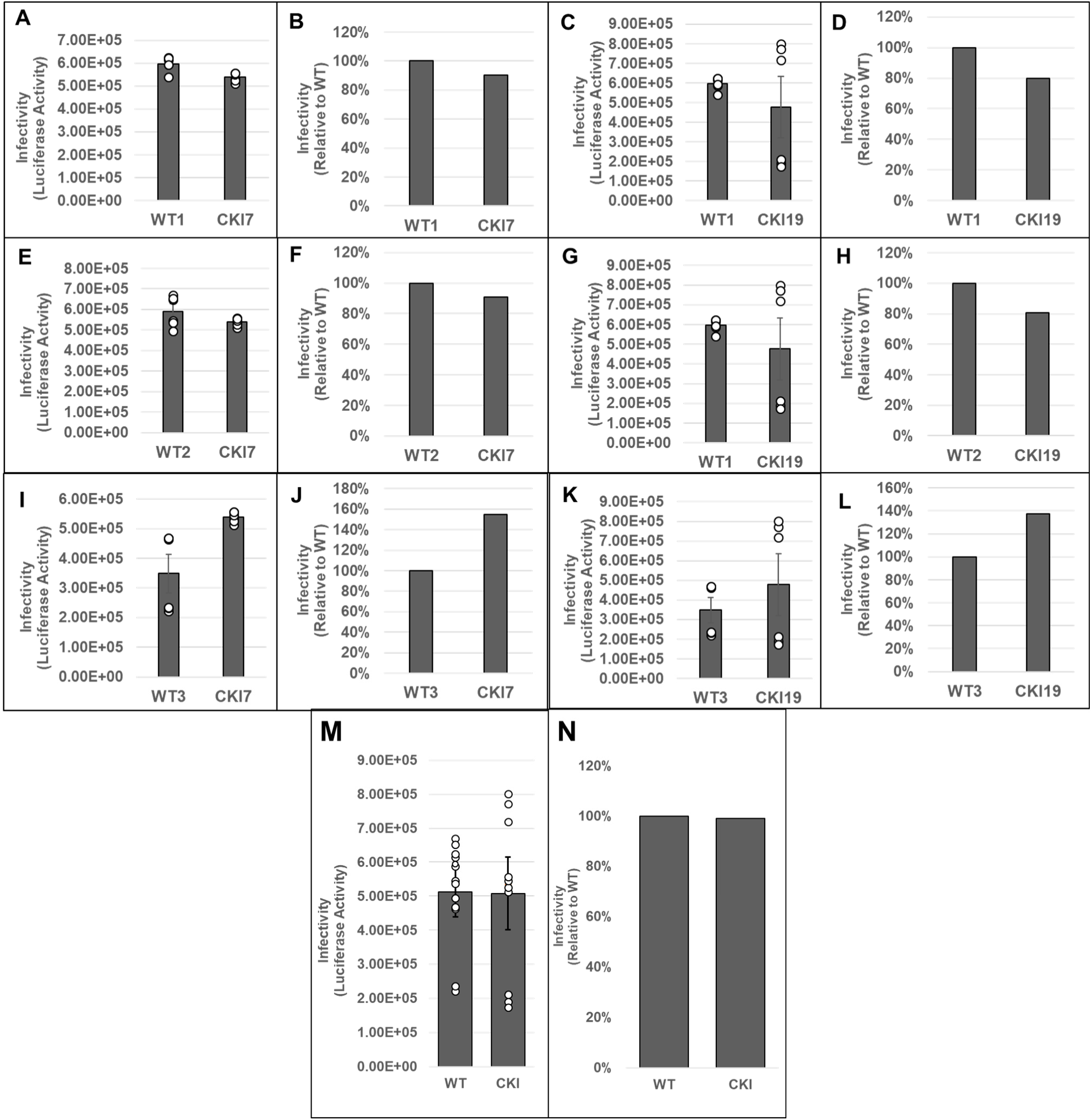
Effects of disrupting the CPSF6-CA interaction on HIV-1 infection at 48 hpi. These data are identical to Figure 8, except that the infection timecourse was for 48 h. **(A, C, E, G, I, and K)** luciferase activity; **(B, D, F, H, J, and L)** Percent infectivity relative to respective control clones. **(M)** Infectivity values sorted by editing treatment (CPSF6 non-edited vs edited). Colored circles individual data point from corresponding cell line. **(N)** Displays aggregate infectivity as a percent relative to WT. Data shown are mean values from three independent experiments, each conducted in triplicate. Error bars represent the standard error of the mean (SEM) and the p-values (*) represents statistical significance (p< 0.05) between the control and CKI cells.

## DISCUSSION

HIV-1 infection is dependent on the integration of PIC-mediated viral DNA integration into gene dense regions of the host genome [65] [120]. Particularly, SPADs and speckle-associated chromatin, which lie in close proximity to nuclear speckles are highly favored targets for HIV-1 integration [117]. A growing body of literature supports that CPSF6 is responsible for targeting the PIC to nuclear speckles for subsequent IN-mediated integration into SPADs [76, 79, 83, 84, 91, 117]. However, several key questions underlying the mechanism by which CPSF6 promotes integration targeting remain unanswered. These include: 1) does the engagement of CPSF6 with HIV-1 CA at a specific location (cytoplasm, NPC, nucleoplasm, or speckles) dictate PIC targeting, 2) are there other viral or host factors that cooperate with CPSF6 for integration targeting, 3) does CPSF6 remain bound to the PIC-associated CA until integration, and 4) importantly, what role does CPSF6 play in PIC function? In this study, we probed the effects of CPSF6 on PIC-mediated viral DNA integration in vitro and in infected cells.

To test a direct role of CPSF6 in HIV-1 integration, we first studied the PIC-the biochemical machinery that carries out viral DNA integration in an infected cell [65, 66]. Due to technical challenges associated with reproducible extraction of PICs from isolated nuclei, we studied PICs extracted from the cytoplasm of infected cells. We found that the PICs from CKO cells exhibited significantly lower integration activity, and calculation of specific PIC activity demonstrated that the lower activity of CKO PICs was not due to lower number of PICs formed in these cells (Fig. 1). Similarly, the integration activity of PICs extracted from CKI SupT1 cells, recoded to disrupt the CPSF6-CA interaction, exhibited significantly lower integration activity when compared to WT SupT1 cell PICs (Fig. 2). Interestingly, the relative reduction in the integration activity of CKI SupT1 PICs was more substantial than the relative reduction in the integration activity of CKO cell PICs. Thus, CPSF6 may exert a dominant effect on PIC function in physiologically-relevant HIV-1 cell types as compared to naturally non-permissive cells, a prediction that requires further investigation. A direct role for CPSF6 in HIV-1 integration was also supported by the data from in vitro integration activity measurements of PICs supplemented with CPSF6 protein (Fig. 3). Addition of CPSF6 protein stimulated the activity of PICs extracted from both CKI and control cells (Fig. 3). However, the extent of increase in integration activity in the presence of CPSF6 was higher for the CKI PICs when compared to the control PICs. Considering the significantly lower integration activity of CKI PICs, supplementation with CPSF6 most likely helps in the partial restoration of the integration activity of these PICs. Indeed, supplementation with 2.0 μM CPSF6 led to ∼7-fold increase in the integration activity of CKI PICs. Conversely, the lack of a stimulation in the activity of control PICs with 2.0 μM CPSF6 suggested a possible saturating effect of CPSF6 on PIC function. These biochemical studies suggest that PIC integration activity is stimulated by CPSF6, and optimal PIC activity depends on the CPSF6-CA interaction.

Our biochemical studies provided in vitro evidence for a direct and functional role of CPSF6 in viral DNA integration by the PIC. Next, we sought to validate the relevance of this observation in HIV-1 infected target cells. For this, we probed CPSF6’s role in HIV-1 integration into the host genome. Our results demonstrated that HIV-1 integration was significantly reduced in CKI cells when compared to WT control cells (Fig. 4A-B). HIV-1 CA mutants incapable of binding to CPSF6 showed no such integration defect in CKI cells when compared to the control cells (Fig. 5), thus confirming that CPSF6’s positive effect on integration is dependent on its binding to the viral capsid. However, HIV-1 integration is dependent on both viral DNA synthesis and viral DNA import into the nucleus [65]. Either of these steps could be influenced by CPSF6-CA binding, since CA regulates both reverse transcription and post-reverse transcription steps [45, 46, 84, 85, 120–123] and the CPSF6-CA interaction affects nuclear entry and post-entry steps of HIV-1 infection [111, 117, 124]. Therefore, we probed whether CPSF6’s effect on HIV-1 integration was dependent on the preceding steps of viral DNA synthesis and nuclear entry.

Notably, viral DNA synthesis was stimulated rather than inhibited in CKI cells (Fig. 4C-D), implying that engagement of CPSF6 with the HIV-1 capsid may negatively impact viral DNA synthesis in T cells. Previous analyses of cytoplasmically mislocalized C-terminal truncation CPSF6 variants reported significant reductions of viral DNA synthesis [76, 80, 125]. Moreover, the higher level of viral DNA synthesis in CKI cells translated to increased HIV-1 nuclear entry (Fig. 4E-F). This was surprising, as CPSF6 depletion was previously reported to yield marginal reductions in HIV-1 nuclear import in certain cell types [79, 109]. Nevertheless, we predict that lack of CPSF6 binding may allow prolonged engagement of CypA with the capsid, since CypA is known to prevent CPSF6-358 mediated restriction of the HIV-1 capsid [85]. Thus, CypA-mediated stabilization of the capsid could plausibly provide a microenvironment suitable for more efficient reverse transcription in CKI cells. In this scenario, higher viral DNA synthesis would generate higher number of 2-LTR circles because only a very small percentage of viral DNA are integrated into the host genome [126]. However, detailed kinetic studies of viral DNA synthesis, 2-LTR circle formation and viral DNA integration in cells depleted of either CypA or CPSF6 or both are required to validate this speculation.

Our studies established that CPSF6 promotes HIV-1 PIC activity, and disruption of CPSF6-CA binding reduces viral DNA integration into the host genome. Importantly, PIC-associated viral DNA is preferentially integrated into active chromatin including highly transcribed genes, gene-dense regions, activating epigenetic marks, and SPADs [84, 120, 127]. Accordingly, integration is disfavored in heterochromatin regions containing repressive epigenetic marks and LADs [82, 127]. One of the primary functions of CPSF6 is to direct the PIC away from the LADs located near the nuclear periphery to promote selective integration into SPADs [76, 79, 83, 84, 91, 117]. So, when CPSF6 is depleted or CPSF6 cannot bind to capsid, the PIC remains closer to the nuclear lamina leading to integration into LADs [54, 79, 82, 109, 128]. Therefore, we asked if, HIV-1 DNA integration targeting was altered in CKI cells due to the loss of the CPSF6-CA interaction. To test this, we mapped HIV-1 integration into genomic features, including genes, SPADs, LADs, CpG islands, TSSs and gene dense regions (Fig. 6). Predictably, HIV-1 integration into genes, gene dense regions and SPADs was significantly enriched in the control cells. In contrast, there was a significant reduction in integration into genes, gene dense regions and SPADs in the CKI cells. Concurrently, the LAD regions served as a major target for viral DNA integration in the CKI cells. This profound integration retargeting is a consequence of the lack of CPSF6 binding to the HIV-1 capsid.

Our PIC studies and cell-based infection assays demonstrated that CPSF6 is important for viral DNA integration. Since viral DNA integration is required for productive infection, we carried out single-round infection studies in both CKO and CKI cells. As previously reported [83], higher HIV-1 infection was observed in HEK293T CKO cells at both 24 hpi (Fig. 7) and 48 hpi (Suppl. Fig. 3). The effects of CPSF6 on HIV-1 infection seem to be cell-type dependent. For example, HIV-1 infection has been reported to be marginally enhanced in CKO HEK293T cells and certain CPSF6-depleted cell lines in single-round infection assays [68, 83, 109, 129, 130]. In contrast, multi-round HIV-1 replication was reduced in CPSF6-depleted primary macrophages [80]. Similarly, depletion of CPSF6 in primary, resting CD4+ T cells reduced HIV-1 infection in single round infection assays [131]. Nevertheless, these prior cell models were instrumental in defining CPSF6’s role during HIV-1 nuclear entry, nuclear trafficking and integration targeting. However, prior cell line models were limited in probing a direct role of CPSF6 in PIC function since CPSF6 regulates multiple steps before integration and each of these steps could affect PIC function. Additionally, prior CKO cell lines, of HEK293T and HeLa cell origin, were sub-optimal to address whether CPSF6’s effect on PIC function was a direct consequence of CA binding or perhaps due to an indirect effect of alterations in cellular physiology associated with depletion of a key cellular protein. To circumvent such limitations, we recoded the critical FG motif in CPSF6 in SupT1 cells (Suppl. Fig. 1-2). Single round infection assays showed significant reduction in viral infectivity in two different CKI cell lines at 24 hpi (Fig. 8). However, by 48 hpi, this infectivity defect had dissipated (Fig. 9). Collectively, these results suggests that the detrimental effect of disrupting the CPSF6-CA interaction on HIV-1 infection is short lived, which is consistent with the notion that CPSF6’s effect on HIV-1 infection is both context and cell-type dependent.

Our results obtained using a combination of genetic, virology, molecular biology, and biochemical approaches indicate a direct role for CPSF6 in HIV-1 PIC function both in vitro and in infected cells. We propose a model for a direct role of CPSF6 in PIC function (Fig. 10). We predict that CPSF6 promotes viral DNA integration function by binding to PIC-associated CA [54, 132, 133]. Speculatively, binding of CPSF6 could negatively affect the complete or sufficient removal of PIC-associated CA, which is likely required for downstream integration. How might CPSF6 regulate complete or sufficient removal of PIC-associated CA? We predict that as the core-associated PIC is trafficked through the nucleoplasm towards nuclear speckles, CPSF6 and CypA cooperatively regulate CA-removal. This is based on evidence that CypA protects the capsid from restriction by cytosolic CPSF6 [134]. However, when CPSF6 binding is disrupted, CypA continues to be associated with the capsid, leading to insufficient removal of PIC-associated CA and consequently impaired integration. This contention is supported by our recent report that CypA prevents integration of certain HIV-1 CA mutants that escape the inhibitory effect of CD8+ cytotoxic T lymphocytes (CTLs) [101]. However, CypA depletion removes the integration block of these CTL-escape mutants, thus CypA can be inhibitory to PIC function under certain conditions. The capsids of these CTL-escape CA mutants retain the ability to bind to CPSF6, implying that CPSF6 binding may negatively affect removal of PIC-associated CA in the mutant viruses. Additionally, the retention of excess CA may reduce PIC function by interfering with the recently reported role of CPSF6 in liquid-liquid phase separation [76, 135–138]. Experimental evidence to support these predictions are warranted to pinpoint the exact mechanism by which CPSF6 regulates HIV-1 PIC function.

**Figure 10.**
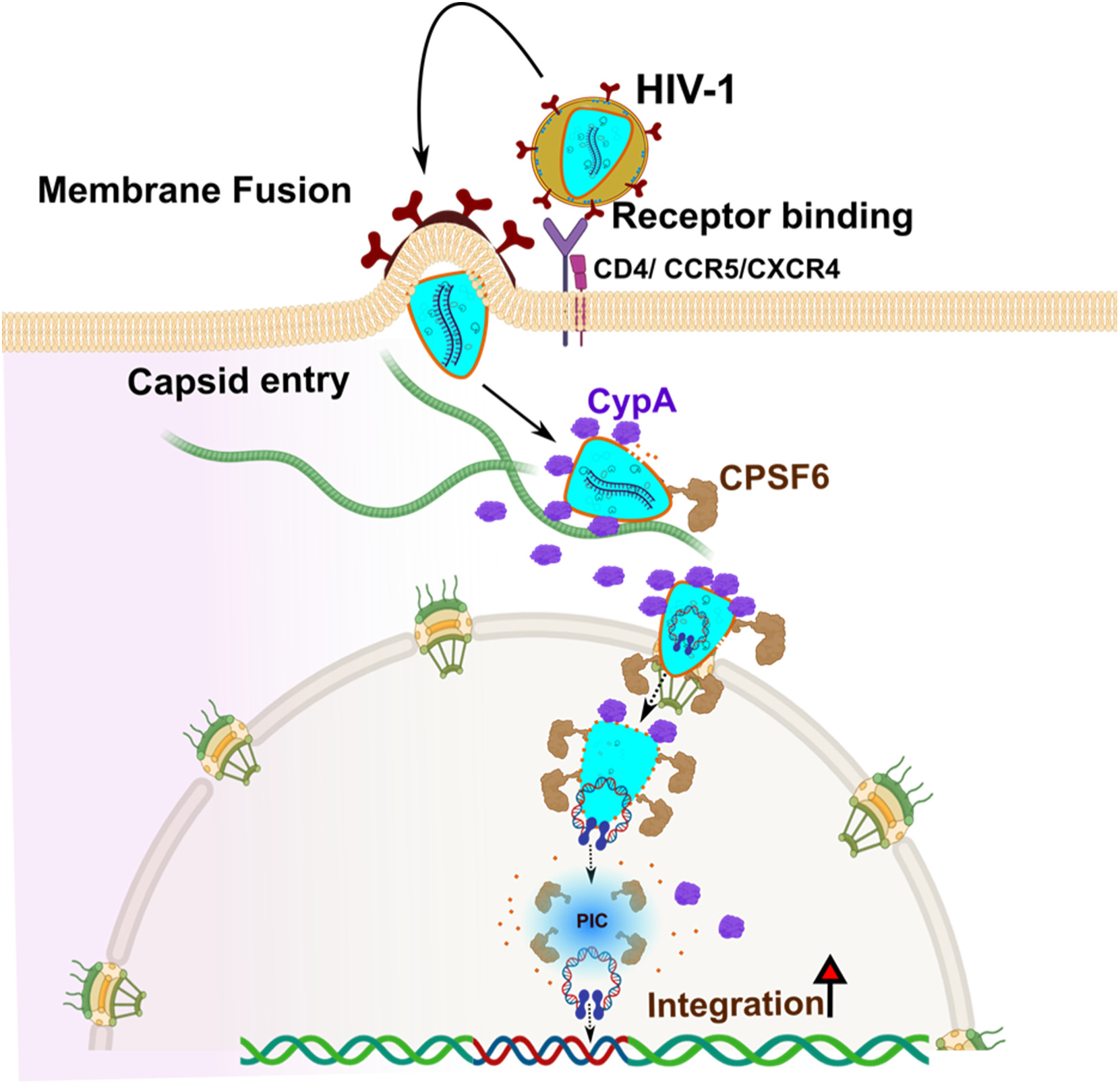
A model supporting the mechanism by which CPSF6 promotes HIV-1 integration. The HIV-1 envelope spike binds to the CD4 receptor and a co-receptor-either CCR5 or CXCR4 to mediate virus entry. HIV-1 capsid is transported via microtubules (shown as green elongated tube) where CypA (deep blue colored structures) binds to the capsid shell. En route to the nucleus, the operationally intact capsid containing partially reverse transcribed viral DNA enters the nucleus through the NPC. Upon nuclear entry, CPSF6 (brown) remains bound to the capsid; the PIC is shown as light blue blob at the site of the integration. Our model predicts that CPSF6 binds to the PIC to promote PIC-mediated viral DNA integration. CPSF6 may influence the final removal of CA (red dots), which would be necessary for IN-mediated viral DNA integration.

## ACKNOWLEDGEMENTS

This work was partly supported by National Institutes of Health grants R01DA057204, R01AI170228, R01AI162694, R01AI136740, R25AI164610 (to CD), R01AI052014, and R37AI039394 (to ANE). This work was also in part supported by the Research Centers in Minority Institutions (RCMI) grant U54MD007586 and the Tennessee CFAR Grant P30AI110527.

## Supplementary Data

**Fig. 1:**
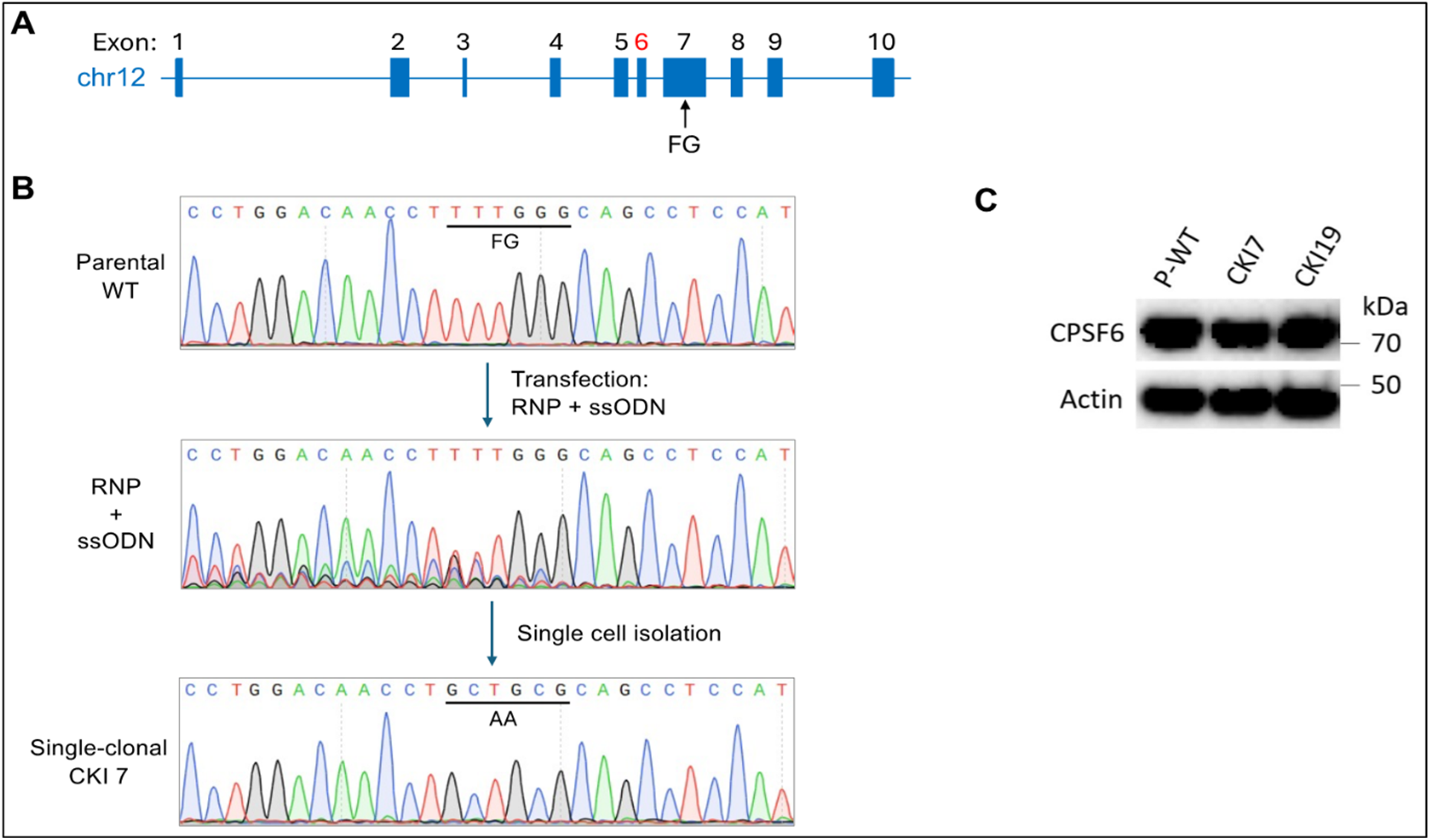
Generation of CKI SupT1 cells by CRISPR-Cas9. **(A)** Schematic of human *CPSF6*; the FG motif is encoded by DNA sequences within exon 7. Exon 6, in red, is most usually removed via alternative splicing. **(B)** Sanger sequencing results of PCR amplicons derived from parental SupT1 cells (upper), transfected cells prior to limited dilution cloning (middle), and the clonally expanded CKI17 cell line. Targeted nucleotides are underlined. **(C)** Immunoblot of CPSF6 expression levels in parental SupT1 cells versus clonally-expanded CKI7 and CKI19 cell lines.

**Fig. 2:**
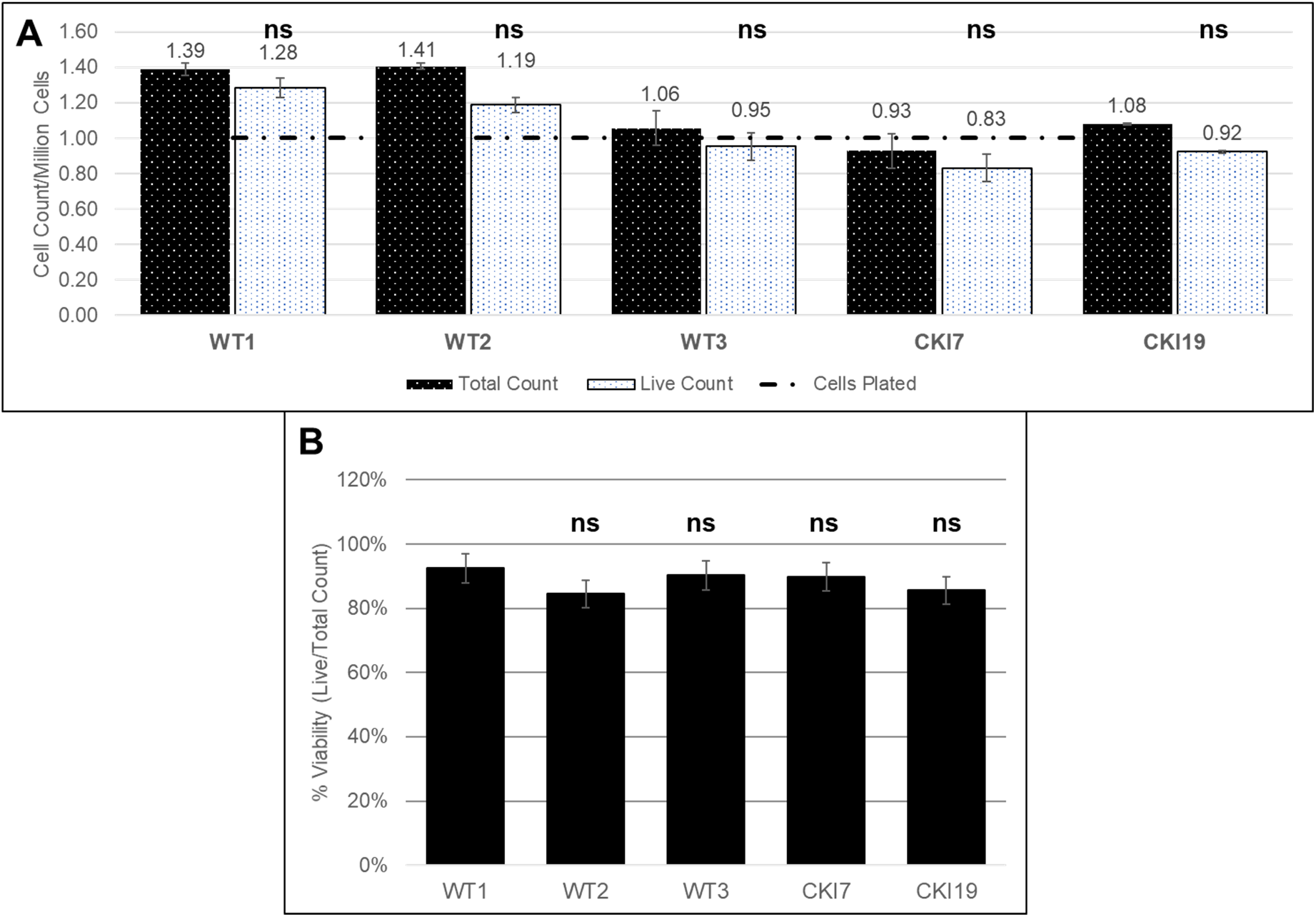
Viability of CKI and WT SupT1 cells. SupT1 cells were compared amongst each other in terms of live and total cell count to assess relative cell viability. Cells (10^6^) were counted prior to seeding in culture plate for 24 h. Following incubation, cells were once again counted and assessed for total cell count, live cell count **(A)** and cell viability **(B)**. Dashed line represents the number of cells seeded prior to seeding. Black patterned bar is representative of an average of 3 total cell counts whereas the white patterned bar is representative of 3 live cell counts. Both values were normalized to 10^6^ cells. **(B)** Shows cell viability (live cell count/total cell count) as a percentage of total cells counted. Error bars are representative of SEM. The p-values (*) represents statistical significance (p< 0.05) compared to WT1. NS represents non-significant difference as compared to WT1.

**Fig. 3.**
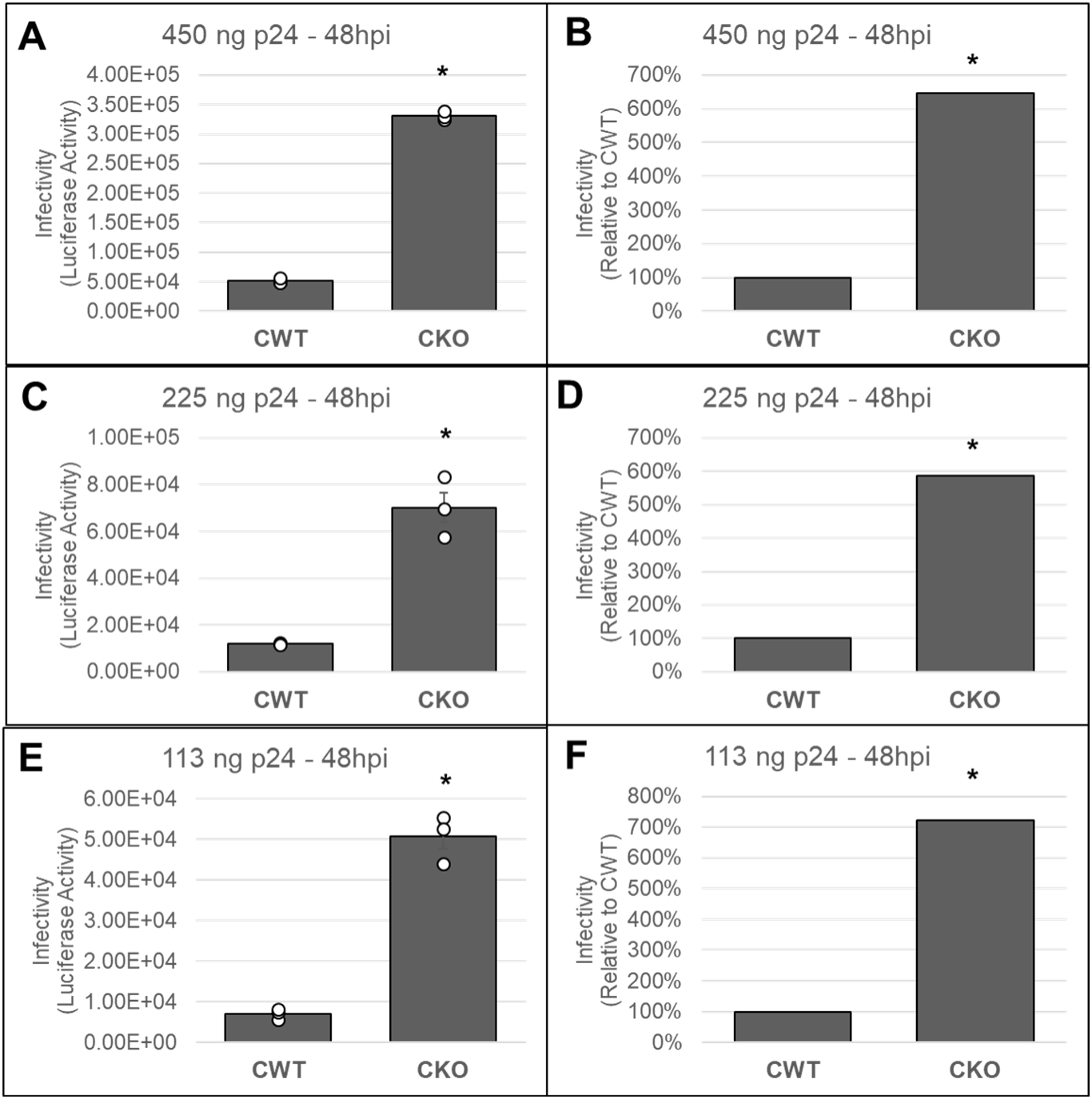
Effects of CPSF6 depletion on HIV-1 infectivity at 48 hpi. HEK293T cells (10^6^) were seeded in 6-well plates for 24 h prior to infection. Following incubation, cell lysates were collected and were processed for assessment of luciferase activity; a surrogate for infectivity. Panel **(A,C,E)** displays the infectivity at 3 concentrations by p24 content of the viral particle: 450 ng **(A)** 225 ng **(B)** 113 ng **(C).** Panels (**B,D,F)** shows the infectivity measurements as a percentage relative to CWT. All experiments were conducted with three independent replicates with 3 technical replicates during the luciferase measurement. Data in **(A,C,E**) are displayed with error bars representing SEM and circles representing replicate data points. The p-values (*) represents statistical significance (p< 0.05) compared to CWT.

**Fig. 4:**
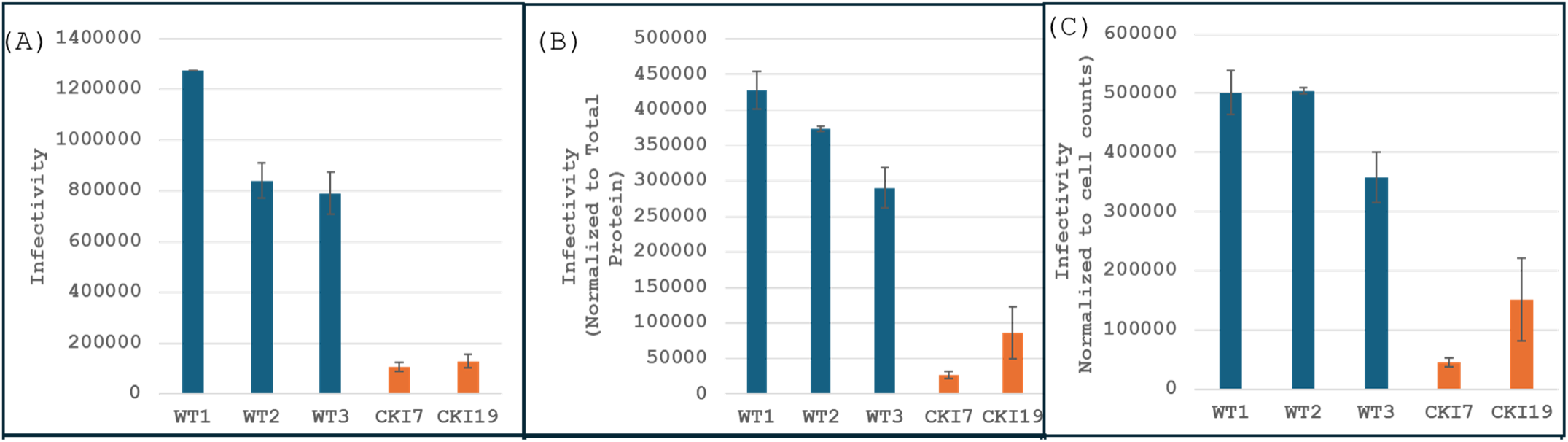
Infectivity in CKI and WT control SupT1 cells. Cells were seeded in 6-well culture plates and inoculated with pseudotyped HIV-1.Luc reporter virus. The entire infection time course spanned 24 h. Subsequently, cell lysates were aliquoted for further processing for total protein (via BCA assay) and luciferase as a surrogate for infectivity. Data shown are representative of mean values from three independent experiments. **(A)** Displays infectivity as represented strictly by luciferase activity. **(B,C)** Infectivity values were normalized to either protein or total cell count respectively. Error bars represent SEM.

